# Transcriptional network analysis of transcriptomic diversity in resident tissue macrophages and dendritic cells in the mouse mononuclear phagocyte system

**DOI:** 10.1101/2020.03.24.002816

**Authors:** Kim M. Summers, Stephen J. Bush, David A. Hume

## Abstract

The mononuclear phagocyte system (MPS) is a family of cells including progenitors, circulating blood monocytes, resident tissue macrophages and dendritic cells (DC) present in every tissue in the body. To test the relationships between markers and transcriptomic diversity in the MPS, we collected from NCBI-GEO >500 quality RNA-seq datasets generated from mouse MPS cells isolated from multiple tissues. The primary data were randomly down-sized to a depth of 10 million reads and requantified. The resulting dataset was clustered using the network analysis tool *Graphia*. A sample-to-sample matrix revealed that MPS populations could be separated based upon tissue of origin. Cells identified as classical DC subsets, cDC1 and cDC2, and lacking *Fcgr1* (CD64), were centrally-located within the MPS cluster and no more distinct than other MPS cell types. A gene-to-gene correlation matrix identified large generic co-expression clusters associated with MPS maturation and innate immune function. Smaller co-expression gene clusters including the transcription factors that drive them showed higher expression within defined isolated cells, including macrophages and DC from specific tissues. They include a cluster containing *Lyve1* that implies a function in endothelial cell homeostasis, a cluster of transcripts enriched in intestinal macrophages and a generic cDC cluster associated with *Ccr7*. However, transcripts encoding many other putative MPS subset markers including *Adgre1, Itgax, Itgam, Clec9a, Cd163, Mertk, Retnla* and *H2-a/e* (class II MHC) clustered idiosyncratically and were not correlated with underlying functions. The data provide no support for the concept of markers of M2 polarization or the specific adaptation of DC to present antigen to T cells. Co-expression of immediate early genes (e.g. *Egr1, Fos, Dusp1*) and inflammatory cytokines and chemokines (*Tnf, Il1b, Ccl3/4*) indicated that all tissue disaggregation protocols activate MPS cells. Tissue-specific expression clusters indicated that all cell isolation procedures also co-purify other unrelated cell types that may interact with MPS cells *in vivo*. Comparative analysis of public RNA-seq and single cell RNA-seq data from the same lung cell populations showed that the extensive heterogeneity implied by the global cluster analysis may be even greater at a single cell level with few markers strongly correlated with each other. This analysis highlights the power of large datasets to identify the diversity of MPS cellular phenotypes, and the limited predictive value of surface markers to define lineages, functions or subpopulations.

## INTRODUCTION

The mononuclear phagocyte system (MPS) [1] is a family of cells present in every tissue in the body including progenitors, circulating blood monocytes and resident tissue macrophages [2-5]. Within each tissue, resident macrophages occupy territories with a regular distribution, commonly associated with epithelial and endothelial surfaces (reviewed in [5]). The proliferation, differentiation and survival of most resident macrophage populations depends upon signals from the macrophage-colony-stimulating factor receptor (CSF1R) initiated by one of two ligands, CSF1 or IL34 [6, 7]. Based upon detection of macrophage-restricted mRNA, including *Csf1r*, the relative abundance of resident macrophages in most organs in mice was shown to reach a maximum in the first week of postnatal life and remains stable thereafter during postnatal growth [8]. Lineage-trace studies in the C57BL/6 strain suggest that many macrophage populations established in the mouse embryo are maintained in adults mainly by self-renewal, whereas others are replaced progressively to differing extents by blood monocytes derived from bone marrow progenitors throughout life [9-11]. Most if not all tissue macrophage populations can be generated and maintained in the absence of blood monocytes due to the intrinsic homeostatic regulation by circulating CSF1 [12]. The precise details of ontogeny, turnover and homeostasis of resident macrophages may not be conserved across mouse strains or species [5]. However, regardless of their steady-state turnover, all resident macrophages including the microglia of the brain can also be rapidly replaced by blood monocytes following experimental depletion ([3-5, 12] and references therein).

Within individual tissues, resident macrophages acquire specific adaptations and gene expression profiles [2, 5, 13-15]. These adaptations contribute to survival as well as function and involve inducible expression of transcription factors and their downstream target genes. At least some of these transcription factors act by regulating *Csf1r* expression. Deletion of a conserved enhancer in the mouse *Csf1r* gene leads to selective loss of some tissue macrophage populations, whereas others express *Csf1r* normally [16]. In the mouse embryo, where abundant macrophage populations are engaged with phagocytosis of apoptotic cells [17], the macrophage transcriptome does not differ greatly between organs. Tissue-specific macrophage adaptation occurs mainly in the postnatal period as the organs themselves exit the proliferative phase and start to acquire adult function [8, 15].

Classical dendritic cells (cDC) are commonly defined functionally on the basis of a proposed unique ability to present antigen to naïve T cells, a concept that requires a clear distinction between DC and macrophages [18]. It remains unclear as to whether cDC should be considered part of the MPS and the extent to which they can be defined by surface markers [12]. The situation is confused by the widespread use of the term DC to describe any antigen-presenting cell (APC) including cells that are clearly derived from blood monocytes [19]. An attempt at consensus proposed an MPS nomenclature classification based upon ontogeny, and secondarily upon location, function and phenotype [20]. The proposal separates monocyte-derived APC from cDC subsets: cDC1, dependent on the transcription factor BATF3, and cDC2, dependent upon IRF4. Some support for this separation came from analysis of an *Ms4a3* reporter transgene, which labelled cells derived from committed granulocyte-macrophage progenitors and distinguished monocyte-derived cells from tissue DC [10]. Secondary classification is based upon cell surface markers that are presumed to be linked in some way to ontogeny. The proposed development pathway of these DC subsets from a common myeloid progenitor, via a common DC progenitor (CDP), has been reviewed recently [21].

Even within tissues resident macrophages are extremely heterogeneous [22, 23]. Since the advent of monoclonal antibodies and later development of transgenic reporter genes [24] numerous markers have been identified that segregate the MPS into subpopulations. Amongst the recent suggestions, LYVE1 was proposed as a marker of macrophages associated with the vasculature [25], CD64 (*Fcgr1*gene) and MERTK as markers that distinguish macrophages from classical DC [26, 27] and CD206 (*Mrc1* gene) as a marker of so-called M2 macrophage polarization [28]. Several surface markers have also been identified that are encoded by genes expressed only in macrophages in specific tissues (e.g. *Clec4f, Tmem119, Siglecf* [22, 23]. Other markers define macrophages in specific locations within a tissue, for example CD169 (encoded by *Siglec1*) in the marginal zone of spleen and hematopoietic islands in bone marrow [29]. In the case of blood monocytes, the subpopulations are clearly a differentiation series in which short-lived LY6C^hi^ “classical” monocytes give rise in a CSF1R-dependent manner [30] to long-lived LY6C^lo^ non-classical monocytes via an intermediate state [11, 30, 31]. This is likely also the case in tissues such as the liver [32] and intestine [33, 34]. More recently, mouse tissue macrophage heterogeneity has been analysed using multiparameter flow cytometry and single cell RNA-seq [35]. Mechanistically, the association between marker expression and cellular function depends upon coordinated transcriptional regulation. One way to identify coregulated sets of transcripts is to cluster large transcriptomic datasets. This approach was used to create transcriptional atlases in multiple species and identify lineage-specific transcription factors and their target genes [36-40]. It enabled the extraction of a generic tumour-associated macrophage signature from multiple large cancer datasets [41]. Previous meta-analysis of large microarray datasets [36, 37, 40] as well as a reanalysis of data from the ImmGen Consortium [42] indicated a clear separation between mouse MPS cells and other leukocyte lineages but did not support the basic premise that markers can separate macrophages from DC or define lineages within the MPS.

Over the past 5 years, RNA sequencing (RNA-seq) has supplanted microarrays as an approach to expression profiling. The recent cascade of interest in tissue-specific macrophage adaptation has produced RNA-seq data for MPS cells isolated from most major organs of C57BL/6 mice. To enable comparative analysis of datasets from multiple laboratories, we devised an automated informatics pipeline employing random sampling of RNA-seq data to a common depth and quantification using the pseudo-aligner Kallisto. Robust transcriptional atlases for the chicken [43] and pig [44] were generated using datasets from numerous divergent sources. Using the same basic pipeline, we identified a total of around 500 RNA-seq libraries generated from isolated macrophage and cDC populations from 24 different studies that sample mouse MPS transcriptional diversity (**Table 1**). Here we apply transcriptional network clustering to this large dataset to analyse transcriptional adaptation across the entire mouse MPS.

**Table 1.** GEO and BioProject accession numbers for samples used in the analysis.

| Accession | BioProject | Reference | Description (markers used) |
| --- | --- | --- | --- |
| GSE125691 | PRJNA<br>517169 | [25] | Interstitial subsets from lung, skin, fat, heart + monocytes and alveolar macs (LYVE1, SiglecF). |
| GSE84586 | PRJNA<br>330530 | [175] | Resident macrophages from heart, kidney, and liver (F4/80, CD11b). |
| GSE94135 | PRJNA<br>369038 | [164] | Three interstitial subsets from lung (Mertk, CD64, CD11b, CD11c, CD206, MHCII) + alveolar macs. |
| GSE95859 | PRJNA<br>378611 | [176] | Brown adipose macrophages (Cx3cr1). |
| GSE114434 | PRJNA<br>471340 | [34] | Monocytes and small intestinal macrophage subsets (CD4, TIM4, CD64) |
| GSE116094 | PRJNA<br>478258 | [152] | Kidney resident and monocyte-derived subpopulations, effect of ischaemia. (F4/80, CD64, CD11c, CD11b, MHCII) |
| GSE122766 | PRJNA<br>506249 | [177] | Brain microglia, bone marrow-derived brain macrophages. (CD45, CD11b, CX3CR1) |
| GSE123021 | PRJNA<br>507265 | [178] | Brain microglia, cortex, cerebellum, hippocampus, striatum (Tmem119) |
| GSE127980 | PRJNA<br>525977 | [131] | Erythroblastic island macrophages from marrow (Epor-EGFP, F4/80, VCAM1, SIGLEC1) |
| GSE135018 | PRJNA<br>557178 | [179] | Alveolar macrophages and peritoneal macrophages, effect of bHLHe40/41 mutation. (SIGLECF, CD11b, CD11c, F4/80). |
| GSE128662 | PRJNA<br>528430 | [32] | Monocyte to Kupffer cell differentiation series (F4/80, CD11b, LY6C, CLEC4F) |
| GSE128781 | PRJNA<br>529096 | [124] | Non-parenchymal brain macrophages, microglia and peritoneal macrophages (MHCII, CD64, CD11b) |
| E-MTAB-6977 | PRJEB<br>27719 | [33] | Macrophage subsets from intestinal lamina propria, serosa and muscularis. (CD64, Cx3cr1 lineage trace) |
| GSE112002 | PRJNA<br>438927 | [130] | Pancreatic islet and peri-islet macrophage populations. Effect of high-fat diet. (F4/80, CD11b, CD11c) |
| GSE103847 | PRJNA<br>407286 | [125] | White adipose and sympathetic neuron-associated macrophages, spleen, microglia. (CD45, Cx3cr1-EGFP, F4/80) |
| GSE68789 | PRJNA<br>283850 | [93] | Mucosal and skin Langerhans cells and DC. (CD103, CD11b, EpCAM, Cd207) |
| GSE128518 | PRJNA<br>527979 | [180] | White adipose macrophages, effect of Trem2 KO. (CD11b, F4/80) |
| GSE107130 | PRJNA<br>419127 | [181] | Brain microglia developmental time course: male and female. Role of microbiome (CD45, CD11b, F4/80, CD64) |
| GSE83222 | PRJNA<br>325288 | [132] | Spleen, intestine, bone marrow macrophages. Effect of engulfment of apoptotic cells. (F4/80, CD11b). |
| GSE95702 | PRJNA<br>378162 | [118] | Monocytes and bone marrow progenitors. CD115, CD135, Ly6C, Cd11b, CD11c |
| GSE130201 | PRJNA<br>534273 | [133] | Dendritic cells, lymph node and spleen. cDC1/cDC2.(CD11c, CD64, MHCII, CD103, Tbx21) |
| GSE120012 | PRJNA<br>491337 | [182] | Cardiac vessel macrophages. MHCII, CCR2, CD64, CD11b |
| GSE140919 | PRJNA<br>519465 | [95] | Monocyte engraftment of colon/ileum. Cx3cr1-EGFP, CD115, LY6C, CD64 |
| GE131751 | PRJNA<br>544681 | [111] | Kidney resident and monocyte-derived macrophage and DC. (F4/80, CD64, CD11c, CD11b, MHCII, Clec9A lineage trace) |

## METHODS

The RNA-seq datasets from within the BioProjects shown in **Table 1** were downloaded from the European Nucleotide Archive (ENA). **Table S1** contains all the SRA and NCBI accessions and sample descriptions. Individual BioProjects differ in methods of mRNA isolation, library preparation and sequencing methods, length, depth and strandedness, but previous analysis in other species [43, 44] indicated that they can still produce comparable expression level estimates. We initially included data from the large ImmGen UL1 project (GSE127267l; GSE124829; see [45]) but this project uses a novel ultra-low input RNA-seq pipeline based upon 1000 sorted cells and the single cell RNA sequencing (scRNA-seq) platform Smartseq2. Our analysis revealed a large batch effect relative to all other samples, and we therefore excluded these data. To reduce possible effects of sampling depth, and generate a common normalisation, the sequences were each randomly down-sampled to a depth of 10 million reads per library as described [43] and requantified using Kallisto v0.44.0 [46]. Kallisto quantifies expression at the transcript level, as transcripts per million (TPM), by building an index of k-mers from a set of reference transcripts and then ‘pseudo-aligning’ reads to it, matching k-mers in the reads to k-mers in the index. The selected BioProjects include subsets of resident tissue macrophages defined using surface markers and isolated by FACS from one tissue as well as temporal profiles of adaptation from monocytes to tissue macrophages. The purpose of this analysis was to identify clusters of transcripts that are robustly correlated. For this purpose, the diversity of transcriptomic space sampled is an asset.

Prior to network analysis, transcripts that were not detected at an arbitrary threshold of 10 TPM in at least one sample were removed to minimise stochastic sampling noise intrinsic in RNA-seq data. Given the nature of the samples, this also helps to reduce the low-level representation of transcripts derived from contaminating cells of non-myeloid origin. Of the 18,175 transcripts that met this minimum threshold, 11,578 were detected in at least 90% of RNA-seq datasets and 6,901 had a median expression >10 TPM. The TPM estimates for the 18,175 transcripts in all of the datasets included are provided in **Table S1**.

Network analysis was performed using the program *Graphia Professional* (https://kajeka.com/graphia-professional/). Pairwise Pearson correlations (*r*) were calculated between all samples to produce a sample-to-sample correlation matrix and inversely between all pairs of genes to produce a gene-to-gene correlation matrix. Gene co-expression networks (GCNs) were generated from the matrix, where nodes represent genes and edges represent correlations between nodes above a defined correlation threshold. For the sample-to-sample analyses an initial screen at the *r* value which entered all samples was performed, followed by subsequent analyses with higher *r* value to remove outliers and reveal more substructure in the networks. For each gene-to-gene analysis the *r* value was adjusted to retain the maximum number of transcripts with the minimum number of edges [43].

## RESULTS AND DISCUSSION

### Expression profiles of individual transcripts

To overview the heterogeneity of the macrophages and the effectiveness of normalisation we first considered the expression profiles of selected individual transcripts to explore the housekeeping genes and surface markers used in studies of MPS cells. The choice of appropriate reference genes for qRT-PCR is a significant issue in many studies, including macrophage differentiation [47]. **Figure 1A** shows candidate house-keeping genes (*Hprt, Actb, B2m, Gapdh, Ppia*) that are commonly used in qRT-PCR as reference genes. These transcripts vary between datasets and BioProjects but in pairwise analysis were only weakly correlated with each other (**Figure 1B**).

**Figure 1.**
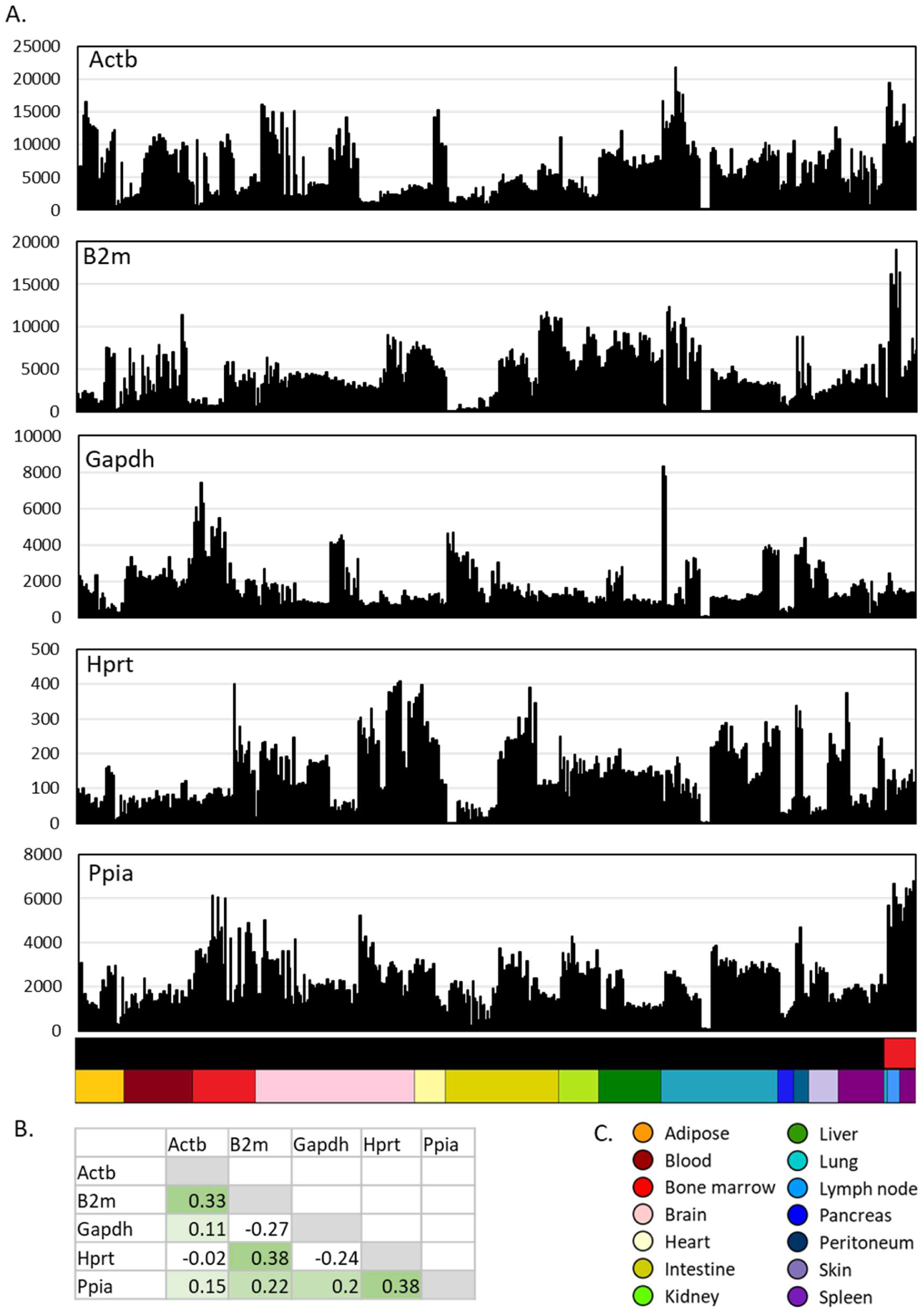
Expression of housekeeping genes across MPS cell populations. **A**. Expression patterns across cells from different tissues. **B**. Correlations (Pearson correlation coefficient) between expression patterns of different housekeeping genes. **C**. Colour code for tissue sources (lower bar, X axis). Upper bar shows cell type: black – macrophage; red – DC.

**Figure 2A** shows the expression pattern of transcripts encoding surface markers used to separate some of the subpopulations herein: *Adgre1* (F4/80), *Cd4, Cd74* (Class II MHC), *Csf1r* (CD115), *Cx3cr1, Fcgr1* (CD64), *Icam2, Itgax* (CD11C), *Lyve1, Mertk, Mrc1* (CD206), and *Tnfrsf11a* (RANK). **Figure 2B** shows a summary of the correlations between them. Consistent with studies using *Csf1r* reporter transgenes [48, 49], *Csf1r* mRNA was universally expressed in MPS cells albeit with significant variation in level, being highest in microglia and lowest in cDC1. *Csf1r* was correlated (r>0.5) with *Adgre1, Fcgr1, Cx3cr1, Mertk* and *Tnfrsf11a* but these transcripts were less correlated with each other. *Mrc1* was reported to be correlated with expression of *Lyve1* and inversely with MHCII [25, 50]. Across the entire spectrum of macrophage transcriptomes, *Mrc1* was correlated with *Lyve1* but was more widely-expressed (**Figure 2A**). There was no evidence of an inverse correlation between *Mrc1* and *Cd74* or other MHCII-associated transcripts.

**Figure 2.**
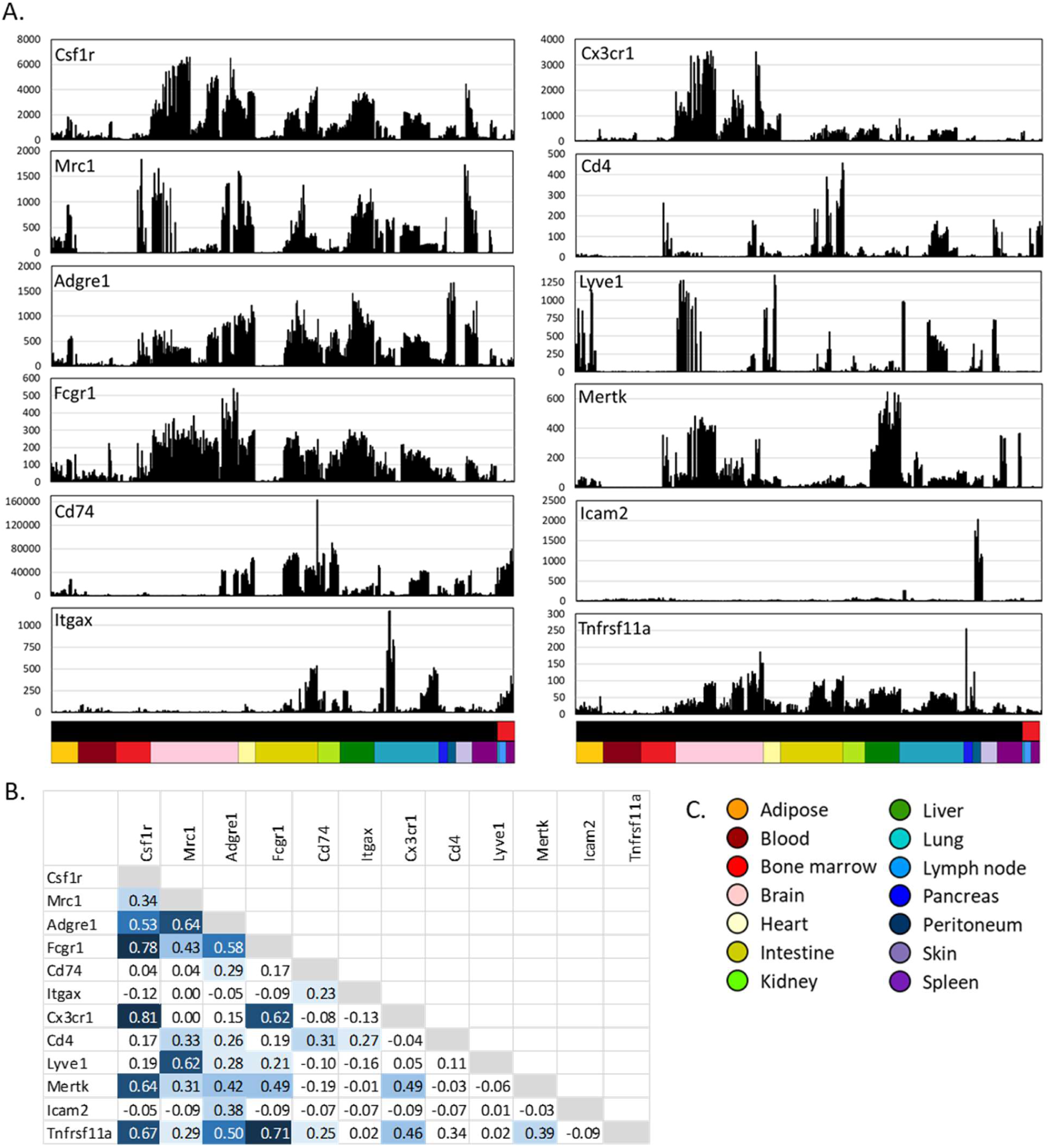
Expression of cell surface marker genes across MPS populations. **A**. Expression patterns across cells from different tissues. **B**. Correlations (Pearson correlation coefficient) between expression patterns of different MPS genes. **C**. Colour code for tissue sources (lower bar, X axis). Upper bar shows cell type: black – macrophage; red – DC.

### Network analysis of relationships of MPS populations and expressed transcripts

To determine whether any transcripts encoding surface markers were correlated with cellular phenotype we used the graph-based network analysis tool *Graphia Professional*. **Figure 3** presents a sample-to-sample correlation matrix generated using the Fruchterman-Rheingold algorithm in *Graphia*, showing the clear segregation of the tissue-specific macrophage populations (**Figure 3A**). Consistent with previous analysis of microarray datasets [37, 39, 40, 42] the isolated spleen, lung and lymph node DC subpopulations clustered together in the middle of the graph (red nodes in **Figure 3B**). Based upon their overall transcriptomic profile, the DC were no more divergent from other MPS populations than the isolated macrophages purified from different tissues were from each other. The apparent relationship to BioProject (**Figure 3C**) occurs mainly because most studies were focussed on a particular tissue or cell type. There may also be impacts from differing methods of extracting and processing RNA and low depth and single end libraries compared to high depth/paired end libraries but these did not produce obvious outliers.

**Figure 3.**
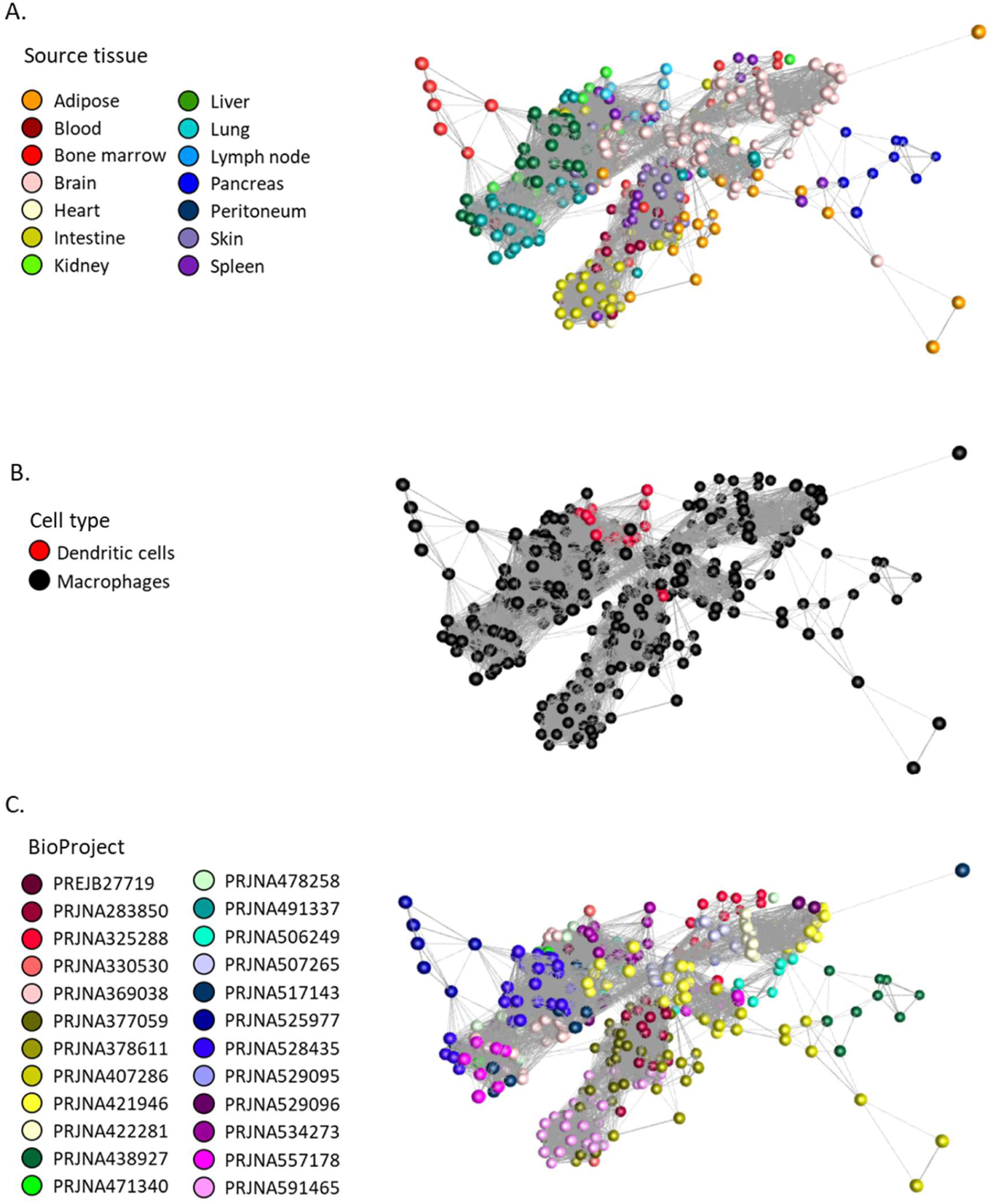
Sample-to-sample network analysis of gene expression in MPS cell populations. Each sphere (node) represents a sample and lines between them (edges) show Pearson correlations between them of ≥ 0.68 (the maximum value that included all 446 samples). **A**. Samples coloured by recorded cell type. **B**. Samples coloured by tissue of origin. **C**. Samples coloured by BioProject.

The gene-centred network (GCN) for the same dataset was developed at an *r* value of 0.75 chosen based on the graph of network size vs correlation threshold shown in **Figure S1. Figure 4A** shows the whole network and **Figure 4B** highlights the tissue specific clusters and those that contain markers of other cell types, as discussed below. **Table S2** summarises the coexpressed gene clusters and the average gene expression profiles of the clusters containing at least 10 nodes (transcripts). The graphs are colour-coded to indicate the tissue origin and cell-type as in **Figure 1** (samples are listed in the Readme sheet of **Table S2**). An additional sheet in **Table S2** provides GO term enrichment of the larger clusters. For ease of visualisation relative to sample information, profiles of surface markers and transcription factors discussed below are provided as an additional sheet in **Table S1. Table 2** provides an overview of the major functional clusters discussed in more detail below. It is beyond the scope of this study to analyse and cite published evidence related to every transcript in detail. In **Table 2**, individual genes from within the cluster have been included based their candidate role as transcriptional regulators and upon known associations with mononuclear phagocyte biology determined by PubMed search on *Genename* AND *macrophage* or *dendritic cell*. On the principal of guilt-by-association [36-40] there are hundreds of other genes within these clusters that have inferred functions in innate immunity and mononuclear phagocyte biology.

**Table 2.**
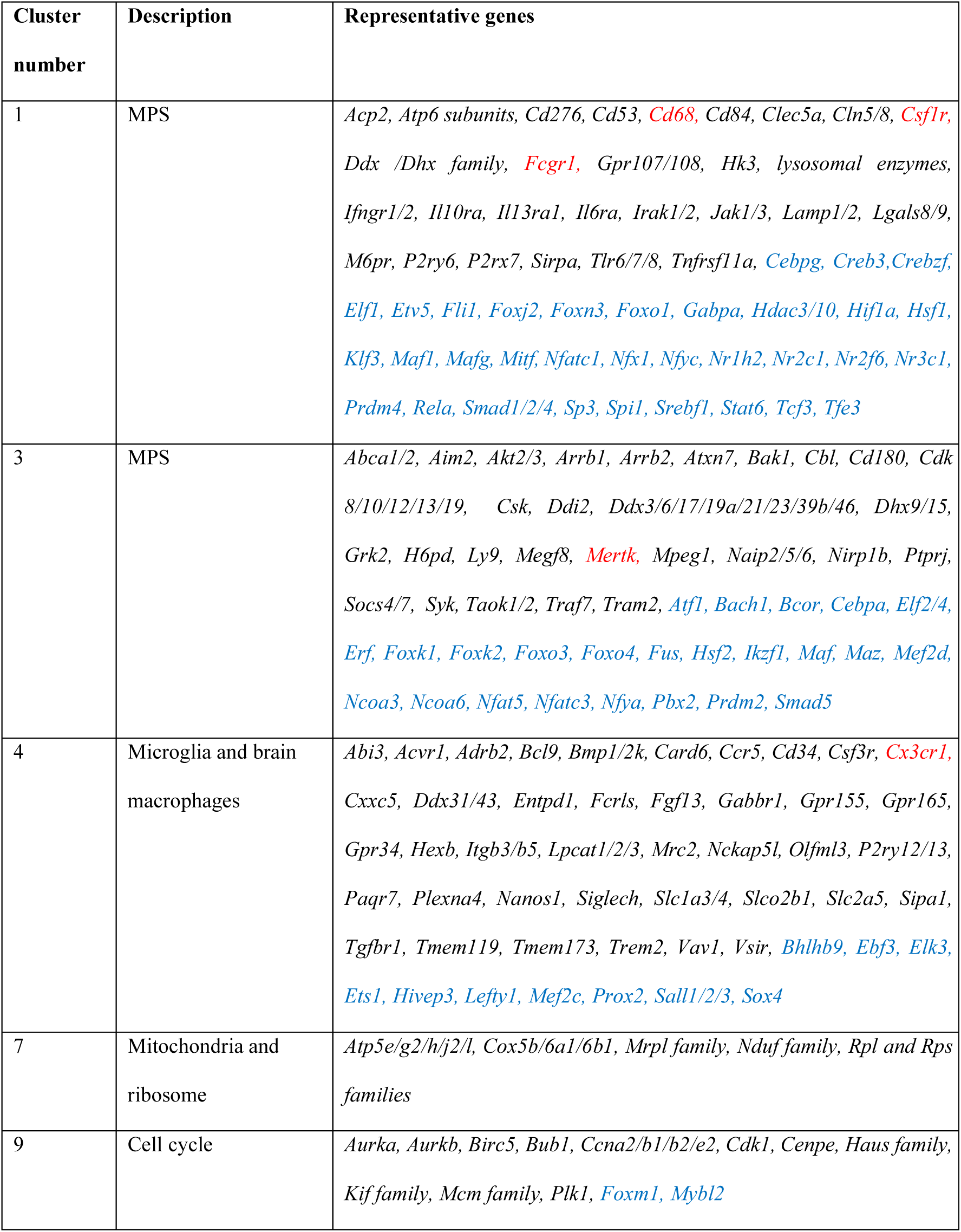
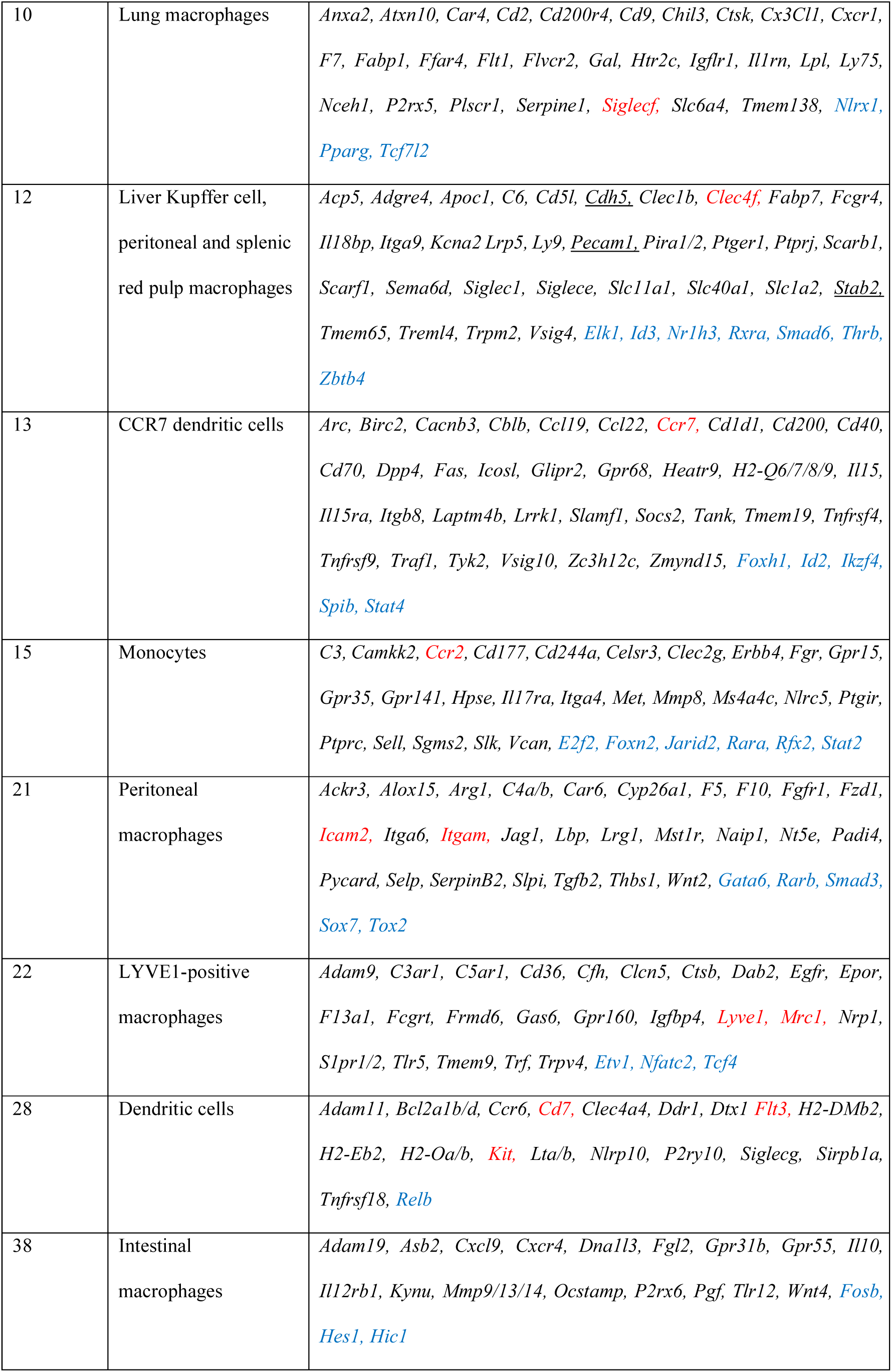
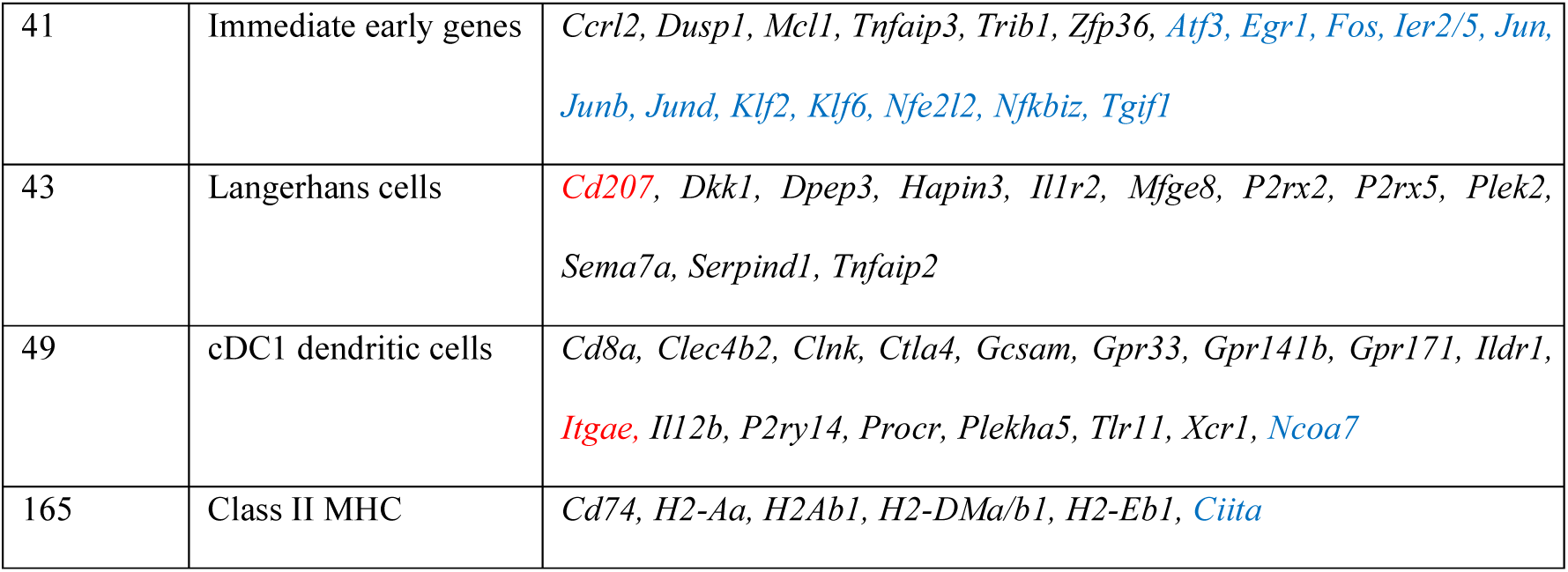
Description of major functional clusters of coexpressed genes in mouse MPS cell samples. Genes in red are key cell surface markers; genes in blue are transcription factors

**Figure 4.**
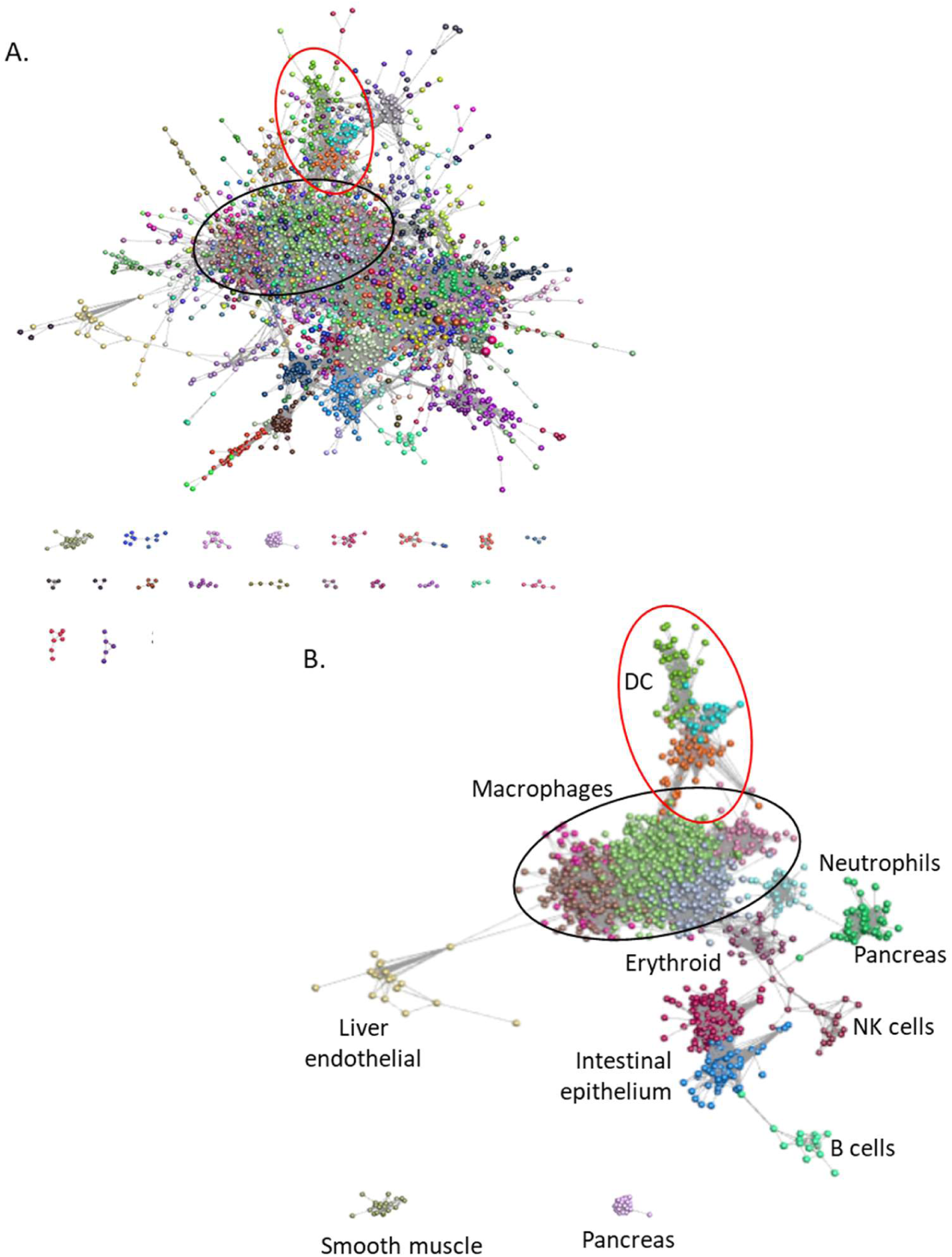
Gene centred network analysis of gene expression in MPS cell populations. Each sphere (node) represents a gene and lines between them (edges) show Pearson correlations between them of ≥ 0.75. Nodes were grouped into clusters with related expression patterns using the MCL algorithm with an inflation value of 1.7. Lists of genes and expression profiles of clusters are presented in **Table** S2. **A**. The network generated by the Graphia analysis. Nodes are coloured by MCL cluster. Lists of genes in all clusters are presented in **Table** S2. Macrophage genes (black oval), DC genes (red oval). **B**. Network showing only major clusters of macrophage genes (black oval), DC genes (red oval) and other cell types.

### Major macrophage-enriched co-regulated clusters

At the chosen *r* value of 0.75, the GCN approach using the normalised data from multiple laboratories identified many co-regulated clusters of transcripts that are consistent with knowledge inferred from smaller datasets.

Cluster 1 is a generic MPS cluster which drives the relatively close association between all of the samples, including the different subclasses of DC, in the sample-to-sample network (**Figure 2A**) and distinguishes MPS cells from other leukocytes. It includes *Csf1r, Fcgr1, Cd68, Sirpa, Tnfrsf11a* and the core myeloid transcription factor gene *Spi1* alongside many other known macrophage-enriched transcription factors [51, 52]. One notable inclusion is the glucocorticoid receptor gene, *Nr3c1*, which mediates transcriptional activation of a wide-range of anti-inflammatory genes in macrophages [53]. As one might expect from the known endocytic and secretory activity of MPS cells, the cluster is enriched for GO terms related to endosome/lysosome and intracellular transport/secretion that are major constitutive functions of mononuclear phagocytes [36] (**Table S2**). Transcripts in Cluster 3 were also expressed widely in MPS cells but the cluster has a distinct average expression profile. Cluster 3 includes genes encoding several forkhead transcription factors (*Foxo3, Foxo4, Foxk1* and *Foxk2*), the key transcriptional regulators of autophagy [54-56], and *Nfat5* which controls macrophage apoptosis [57]. This cluster also contains *Mertk*, the perforin-like immune effector gene *Mpeg1, Aim2*, which encodes a sensor for cytoplasmic DNA [58] and transcripts for numerous DEAH- and DEAD box helicases all implicated in DNA sensing in innate immunity [59]. There are also members of the NAIP family of inflammasome regulators (*Naip2, 5, 6*); reviewed in [60]). We infer that this cluster of transcripts reflects an independently-regulated capacity for innate immune recognition of internalised pathogens. Other than *Mertk*, there is no obvious plasma membrane marker associated with this set of candidate innate immune effector genes.

Genes in Cluster 4 were strongly expressed in samples from brain and include microglia-enriched markers that are depleted in brains of *Csf1r*-deficient mice and rats, such as *Cx3cr1, Tmem119, P2ry12* and the key transcription factor genes *Sall1, Sall2* and *Sall3* [16, 61]. Cluster 9 contains the S phase transcription factor gene *Foxm1* and numerous cell cycle-associated transcripts [62] and the GO term enrichment supports a cell cycle association. The cell cycle cluster was expressed in all isolated MPS populations at various levels consistent with evidence that they are capable of self-renewal in the steady state [5, 12]. The separation of this cluster indicates that proliferative activity is not tightly-linked to any MPS differentiation state or surface marker.

### Identification of a capillary-associated expression cluster

Most macrophages and DC included in this analysis were purified by FACS based upon their expression of specific markers including those shown in **Figure 1B** (see **Table 1**). Chakarov et al. [25] identified a population of pericapillary cells in the lung that expressed LYVE1 and extended their analysis to FACS-separated cells from fat, heart and dermis. Their RNA-seq results are included in our dataset. Based upon analysis of differentially expressed genes, the authors identified a set of genes with high expression in sorted LYVE1^hi^ macrophages relative to LYVE1^lo^ macrophages across the four tissues, including *Mrc1,Timd4, Cd5l, Fcna* and *Vsig4* [25, 50]. The GCN reveals that there is, indeed, a set of transcripts (Cluster 22) that is strongly correlated with *Lyve1* expression across a larger spectrum of tissues. The cluster includes *Mrc1* but excludes *Timd4, Cd5l, Fcna*, and *Vsig4*, which were associated with distinct tissue-specific clusters (**Table 2**). The correlation between *Lyve1* and *Mrc1* is actually lower than the cluster threshold of 0.75 (*r=*0.62, **Figure 2B**). The two genes were included in Cluster 22 because of shared links to other genes. In fact, *Mrc1* was only marginally-enriched in the purified LYVE1^hi^ macrophages from fat, heart, lung and skin [25] and it was highly-expressed in isolated cells from adipose, brain, intestine, kidney and liver that lack *Lyve1* mRNA (see **Table S1**/selected transcripts). We conclude that most LYVE1^hi^ macrophages express *Mrc1*, but the reciprocal relationship does not hold.

The set of co-expressed genes in Cluster 22 suggests a function for LYVE1^hi^ macrophages in control of endothelial biology and vascular permeability. It includes genes for two of the sphingosine-1-phosphate receptors (*S1pr1/2*) which have been implicated in many aspects of inflammation, lymphangiogenesis and angiogenesis [63, 64], the vanilloid receptor (*Trpv4*), which controls capillary barrier function and inflammation [65, 66] and neuropilin 1 (*Nrp1*) which controls endothelial homeostasis [67]. Cluster 22 also contains the erythropoietin receptor gene (*Epor*), which was shown to synergise with S1P to promote apoptotic cell clearance by macrophages [68] and the EGF receptor gene (*Egfr*) which has also been shown to regulate macrophage function in a range of inflammatory models [69]. Indeed, the co-expressed genes might support the known functional association of macrophages with lymphatic as well as blood vessels [70]. The *Lyve1*-associated cluster contains genes for three novel candidate transcriptional regulators *Etv1, Nfatc2* and *Tcf4. Etv1* expression in macrophages has been implicated in functional polarization *in vitro* and the response to altered mitochondrial membrane potential [71]. *Nfatc2* is required for osteoclast differentiation *in vitro* [72] but roles in macrophage differentiation/function have not been explored. *Tcf4* encodes a transcription effector of Wnt/β-catenin pathway, implicated in responses to E-cadherin and other effectors in macrophage differentiation [73].

*Mrc1* is commonly referred to as a marker for alternative or M2 macrophage polarization [74]. Another putative marker of M2 polarization is the somatic growth factor insulin-like growth factor 1 (*Igf1* gene) [75]. *Igf1* was correlated with *Mrc1* (*r=*0.67) but did not form part of a co-expression cluster. It was absent from monocytes and DC but was highly-expressed in most resident tissue macrophages (see **Table S1**/selected transcripts). *Igf1* is CSF1-inducible and of particular interest because of the profound impact of *Csf1r* mutations in multiple species on postnatal growth and development [7]. Unlike hepatocytes and mesenchymal cells, tissue macrophages did not express transcripts encoding the growth hormone receptor (*Ghr*), *Igf1r*, or the *Igf1* binding protein genes (*Igfals, Igfbp1,2,3,5,6*). The exception is *Igfbp4* which was highly-expressed in most macrophage populations and did form part of the *Lyve1/Mrc1*-associated Cluster 22. Interestingly, *Igfbp4* knockout in mice mimics impacts of *Csf1r* deficiency on somatic growth and adipose formation [76, 77].

The intimate association of macrophages with capillaries was evident from the first localization of the F4/80 antigen [78]. *Adgre1* expression was also correlated with *Mrc1* (*r*=0.64; Figure 2B), but it was more widely-expressed than either *Mrc1* or *Lyve1* and therefore not within Cluster 22. *Adgre1* was not enriched in any of the purified LYVE1^hi^ macrophage populations relative to LYVE1^lo^ cells from the same tissue [25]. It was high in most isolated tissue macrophages and induced during differentiation of monocytes *in situ* as in the liver dataset [32] and the intestinal developmental series [33, 34]. F4/80 was proposed as a marker of macrophages of embryonic origin [79] but *Adgre1* was equally high in intestinal macrophages, which turn over rapidly from monocytes [80, 81] and in cDC2. It was also strongly induced during monocyte differentiation to occupy a vacant Kupffer cell niche [32]. Whatever the association with ontogeny, the pattern is rodent-specific. *Adgre1* is a rapidly-evolving gene and the expression pattern also varies across species [82].

### Tissue-specific macrophage clusters

Several co-expressed clusters were associated with MPS cells isolated from a single tissue. Aside from the large brain-enriched expression cluster (Cluster 4) that contains many microglia markers, Cluster 10 was lung-enriched and contains the alveolar macrophage marker *Siglecf* and key transcription factor *Pparg* [15]. Cluster 12 was shared amongst liver Kupffer cells (KC), peritoneal macrophages and splenic macrophages and includes the transcription factors *Id3, Nr1h3* and *Smad6* and markers *Cd5l, Clec4f* and *Vsig4* [15, 32, 34]. A novel finding was the strong coexpression (*r* = 0.81) between *Nr1h3* and *Rxra*, the gene encoding its promiscuous heterodimerisation partner, which is also implicated in control of KC lipid and iron metabolism [83] and may have independent function in innate immune regulation [84].

The average expression of Cluster 12 increased progressively in the monocyte-KC differentiation series [32] included in this dataset (see profile in **Figure S2**). Cluster 12 also reveals the regulated and coordinated expression of the thyroid hormone receptor (*Thrb*), likely mediating the many impacts of thyroid hormones in innate immune function [85]. One other novel candidate regulator identified in this cluster is *Zbtb4* which encodes an epigenetic regulator with a high affinity for methylated CpG. *Zbtb4*^-/-^ mice are viable and fertile but growth retarded compared to littermates [86]. Impacts on myeloid differentiation have not been reported. The transcription factor SPIC is implicated in splenic red pulp macrophage differentiation and iron homeostasis [87, 88]. Although *Spic* mRNA was highest in red pulp macrophages, KC and bone marrow macrophages, it was detected in other macrophage and DC populations and therefore has a unique expression profile. Cluster 21 contains transcripts most highly-expressed in resident peritoneal macrophages and includes the genes for the transcription factor *Gata6* and the retinoic acid receptor (*Rarb*) which control peritoneal macrophage survival and adaptation [89, 90]. The data confirm the specific high expression of the enigmatic plasminogen activator inhibitor encoded by *Serpinb2* in resident peritoneal macrophages, first described >20 years ago [91] and still seeking a function [92].

Genes in Cluster 15, including the monocyte-specific chemotactic receptor *Ccr2*, were highly expressed in classical monocytes. Genes in Cluster 43 were expressed specifically in Langerhans cell (LC). They include the marker *Cd207* (langerin), used in the purification of LC [93] but also expressed at lower levels in many other tissue macrophage populations. This cluster did not include the gene for another LC marker, *Epcam* [93]. It was highly-expressed in LC but also detected in one set of intestinal macrophage samples, most likely a contamination with epithelial cells (Cluster 5, see below). Epidermal LC have at times been considered as DC-like because of their migratory and APC properties but are now considered to be specialized resident tissue macrophages [94]. Unlike most classical DC in lymphoid tissue, they are clearly CSF1R-dependent and share with several other macrophage populations dependence on the conserved enhancer in the *Csf1r* locus [16]. Cluster 43 did not include a transcriptional regulator specific to LC. In common with several other macrophage populations, LC differentiation is regulated by TGFβ signaling, involving transcription factors RUNX3 and ID2 [94]. Both transcription factor genes were highly-expressed in LC but also present in several other tissue macrophage populations.

Intestinal macrophage-enriched gene expression profiles, which have not previously been identified, emerge in Cluster 38. Two large separate datasets of intestinal macrophages were included here [33, 34], both likely reflecting a differentiation series of adaptation from blood monocytes to resident intestinal tissue macrophages [5]. In one case, CD4 and TIM4 were used as markers [34] but each of these markers is shared with other macrophage populations. *Cd4* mRNA expression was shared uniquely with lung, skin and kidney macrophage subpopulations (see **Figure 2B**). A third dataset tracks the adaptation of transferred blood monocytes to the intestinal niche [95]. Cluster 38 identified *Cxcr4* as a candidate intestinal macrophage marker consistent with their continuous derivation from CXCR4^+^ monocytes. The high expression of *Wnt4* in lamina propria macrophages was recently confirmed by IHC. Conditional deletion of *Wnt4* using *Itgax*-cre led to dysregulation of immunity against an intestinal parasite [96]. WNT4 is a candidate mediator of the key trophic role of lamina propria macrophages in the intestinal stem cell niche [97]. *Fosb, Hes1* and *Hic1* encode identified potential transcriptional regulators of intestinal macrophage differentiation and adaptation. HES1 inhibits inflammatory responses in macrophages and contributes to gut homeostasis [98, 99]. FOSB has not previously been implicated in macrophage adaptation to any niche. Unfortunately, we were not able to include data from a microarray analysis of resident colonic macrophages which identified a set of 108 genes >2-fold higher in the colon relative to other macrophage populations in the ImmGen database [100]. However, Cluster 38 confirmed the gut macrophage-specific expression of several of these transcripts including *Dna1l3, Fgl2, Gpr31b, Hes1, Mmp13, Ocstamp, Pgf* and *Tlr12*.

There were no unique expression profiles enriched in macrophages isolated from any other major tissues including adipose, brain (non-microglia), heart, kidney, pancreas or skin. The abundant resident macrophages of adipose are especially topical in light of the obesity epidemic. The literature on adipose macrophages focusses on “M2-like” markers [101]. Amongst resident macrophage populations, *Apoe* and *Retnla*, both detected in most tissue macrophages and not included in a cluster, were highest in adipose-derived macrophages. RETNLA (aka RELMα) has been referred to as an adipokine, regulated by food intake and controlling lipid homeostasis [102]. Kumamoto *et al*. [103] claimed that *Retlna* was co-expressed with *Mgl2* (another putative M2 marker) in many mouse tissues including adipose and attributed it a role in maintenance of energy balance. The two transcripts were not correlated in this larger dataset. In fact, *Mgl2* was part of a small cluster (Cluster 83) with *Cd14*. Like *Retnla*, mRNA for the related lectin, MGL1 (*Clec10a* gene), also considered an M2 macrophage marker [101] was highest in the adipose-associated macrophages but also expressed in macrophages from other tissues including dura, heart, lung and skin (Cluster 101).

### Dendritic cell co-expression clusters

Despite evidence that it is expressed by many resident tissue macrophages (reviewed in [24]), CD11C (ITGAX) is still widely-used as a surface marker in mouse DC purification. Ongoing studies of the impacts of conditional mutations using *Itgax-*cre continue to be interpreted solely in terms of DC specificity (e.g. [96, 104, 105]. Consistent with the literature, *Itgax* was expressed in multiple macrophage populations (**Figure 2A**) at levels at least as high as in purified DC, and correlated only with *Cd22, Cd274* (encoding PD-L1), *Csf2rb, Csf2rb2, Slc15a3, Tmem132a* and the transcription factor gene *Prdm1* (Cluster145). Class II MHC is also commonly used as a marker to purify DC and expression is obviously a prerequisite for antigen presentation to T cells. The ImmGen consortium compared DC from multiple sources with various macrophage populations to identify transcripts that distinguish DC from macrophages [26, 27]. Since the macrophages used for comparison were MHCII^lo^, the DC signature included class II MHC genes. In our meta-analysis, one small cluster (Cluster 165) contained the transcription factor gene *Ciita* and its targets *Cd74, H2-Aa, H2Ab1, H2-DMa/b1, H2-Eb1*. The genes in this cluster were clearly highly-expressed in many tissue macrophages, (see profile for *Cd74* in **Figure 2A**) but regulated independently of any other markers and expressed no higher in cells annotated as DC than in cells annotated as macrophages from intestine, lung, heart and kidney. Interestingly, again highlighting the issue with a definition of DC based upon unique APC function, isolated lung MHCII^hi^ interstitial macrophages were as active as cDC2 in antigen-presentation assays *in vitro* [25].

The GCN analysis did identify three separate DC-associated co-expression clusters that are consistent with current knowledge of putative DC subsets and adaptation in mice [20, 21, 106]. Cluster 13 includes *Ccr7* and transcription factors *Spib* and *Stat4*, Cluster 28 includes *Flt3, Kit* and the transcription factor *Relb* and Cluster 49 includes cDC1 markers *Itgae* (CD103) and *Xcr1*. CCR7 is associated with DC migration [107] and the transcript was abundant in both cDC1 and cDC2 isolated from spleen and lymph node (LN). By contrast, the expression was much lower in isolated lung DC and in kidney DC from a separate dataset (see below), similar to levels in isolated macrophages from multiple tissues. Several putative DC markers were excluded from DC-specific clusters. The transcription factor gene *Batf3*, implicated in cDC1 differentiation [108] did not form part of a cluster and was detected in most macrophage populations (consistent with[15]). Similarly, IRF4 has been attributed a specific function in cDC2 differentiation [105]. *Irf4* mRNA was more abundant in cDC2 compared to cDC1 but was also expressed in monocytes and monocyte-derived macrophage populations. Transcripts encoding NFIL3 and IRF8, which interact in the regulation of cDC1 differentiation [109] were also highly-expressed in cDC2 and in monocytes and many tissue macrophages. Although the transcription repressor gene *Zbtb46*, encoding a putative DC lineage marker [110] was highest in DC it was also detectable in most isolated tissue macrophages, notably in kidney and lung. Another putative DC marker gene, *Clec9a* [111] also clustered independently because of expression in isolated intestine, kidney, liver and lung macrophages.

Interestingly, tissue macrophages may contribute to homeostatic regulation of cDC differentiation. The transcript encoding the FLT3 ligand (*Flt3l*) was expressed constitutively to varying degrees in all of the MPS populations studied. Fujita *et al*. [104] showed that FLT3L is cleaved from the cell surface of expressing cells by ADAM10. Conditional deletion of *Adam10* using *Itgax-*cre led to reduced differentiation of cDC2. *Adam10* is also expressed by CD11C^+^ macrophages; it forms part of Cluster 3, low in monocytes and expressed by all resident macrophages at higher levels than in DC.

Aside from CLEC9A, many other lectin-like receptors have been proposed as DC markers and inferred to have a function in antigen uptake. **Figure 5** shows the profiles for the 12 members of the so-called dendritic cell immunoreceptor (DCIR) family. The original member of this family, *Clec4a2*, the likely ortholog of the single *CLEC4A* gene in humans, encodes a lectin with a broad binding specificity for mannose and fucose [112]. Studies on knockout mice lacking *Clec4a2* continue to be based upon the claim that the lectin is mainly expressed by DC [113] but the global analysis showed that it is more highly-expressed in most isolated macrophage populations. Two of the DC-associated clusters contained other members of the family, *Clec4a4* and *Clec4b2. Clec4a4* has been attributed a specific role in cDC1 dendritic cell function [114] but it was equally expressed in cDC2 and forms part of Cluster 28. Most of the Clec4 genes in the mouse genome are in a single location on Chromosome 6. They also include macrophage-inducible C-type lectin (Mincle) encoded by *Clec4e*, which mediates innate immune responses to *Candida* [115]. The related *Clec4f* (Kupffer cell marker) and *Cd207* (langerin) are located together in a separate locus on Chromosome 6. Each of the *Clec4* genes had a unique expression profile in tissue macrophage populations. Analysis of the entire dataset reveals that “DCIR” is a misnomer for this family. The DC designation has also been misapplied to other putative markers, including DC-SIGN (*CD209* in humans), DEC205 (*Ly75*) and DC-HIL (*Gpnmb*). In mice there are multiple *Cd209* paralogs. *Cd209b* was highly-expressed in marginal zone macrophage populations in spleen and is *Csf1r*-dependent [61]. These cells have not been successfully isolated by tissue disaggregation. Four members of the CD209 family (*Cd209a,d,f,g*) were co-expressed in a unique pattern (Cluster 100) together with *Cbr2, Ccl24* and *Clec10a. Ly75* was detected in both cDC subpopulations but was most highly-expressed in lung macrophages (Cluster 10).

**Figure 5.**
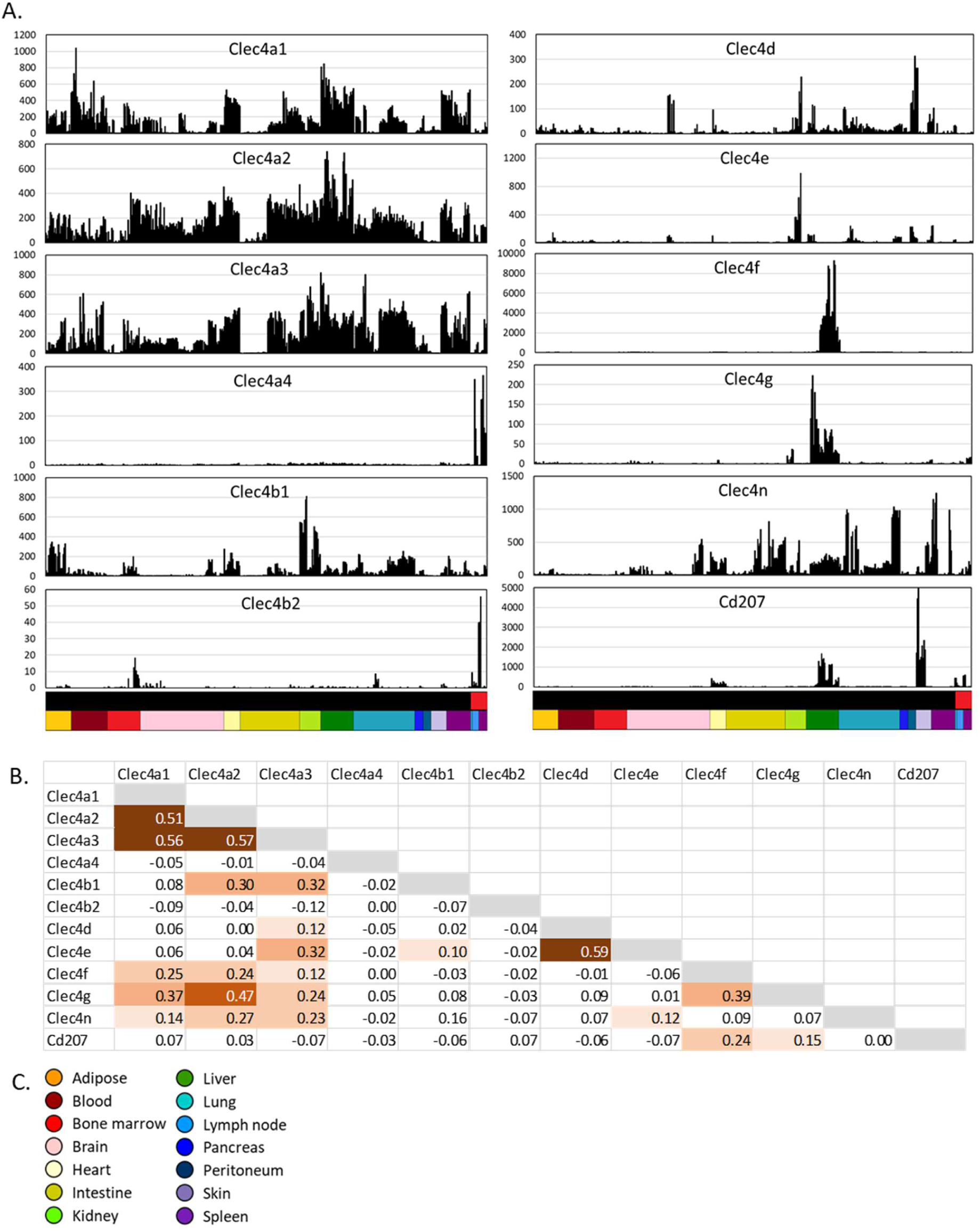
Expression of members of the dendritic cell immunoreceptor (Clec4) family across MPS cell populations. **A**. Expression patterns across cells from different tissues. **B**. Correlations (Pearson correlation coefficient) between expression patterns of different Clec4 genes. **C**. Colour code for tissue sources (lower bar, X axis). Upper bar shows cell type: black – macrophage; red – DC.

CD64 was used as an exclusion criterion to remove or separate macrophages from DC or to enrich macrophages in all of the datasets included herein based upon the earlier studies of the ImmGen Consortium [26]. This exclusion was clearly successful in that all the purified DC have very low *Fcgr1* (**Figure 2A** and **Table S1**), but the expression of this gene in macrophage populations was also highly variable. As a simple screen for additional markers that distinguish all “macrophages” from all “DC”, we averaged expression across all macrophage and DC samples and compared them (see **Table S1**). Amongst the transcripts that were robustly-expressed and highly-enriched in macrophages to at least the same extent as *Fcgr1*, those encoding surface markers were also variably-expressed amongst macrophage populations. However, we identified three transcription factor genes, *Cebpb, Mafb* and *Klf10*, that were apparently excluded from all of the cDC. The role of *Cebpb* in macrophage differentiation is well-recognised [116-118] and one of the datasets includes progenitors from *Cebpb*^-/-^ mice [118]. There is evidence of a negative feedback relationship with *Irf8* in monocyte-derived DC [119]. *Cebpb* was detected in most tissue macrophages but uniquely excluded from some populations, notably the heart and intestinal muscularis. *Mafb* has been proposed previously as a lineage marker separating macrophages from DC [120, 121]. The literature on *Klf10* is more limited, with evidence that it participates in TGFβ-induced macrophage differentiation [122].

### Resident macrophage activation during isolation

Cluster 41 contains numerous immediate early genes (IEG) encoding transcription factors and feedback regulators (e.g. *Fos, Egr1, Dusp1*) consistent with evidence that isolation of cells from tissues produces cell activation, from single cell sequencing of disaggregated cells [123, 124]. In many samples, IEG were amongst the most highly-expressed transcripts. The majority of isolated macrophage populations also had high levels of macrophage-specific LPS-inducible genes. Cluster 224 contains *Ccl2, Ccl7, Ccl12, Cxcl1* and *Il6*, Cluster 329 includes *Il1b* and *Ptgs2* (Cox2), and Cluster 485 contains *Tnf* and inducible chemokines *Ccl3* and *Ccl4*. The anti-inflammatory cytokine *Il10*, which is also LPS-inducible, formed part of the intestinal macrophage cluster (Cluster 38). IL10 is essential to intestinal homeostasis [80] but *Il10* mRNA was detected in only one of the three intestinal macrophage datasets [34] alongside very high expression of IEG and proinflammatory cytokines (e.g. *IL1b, Tnf*). Inflammation-associated transcripts were highlighted as evidence of activation *in vivo* in sensory neuron-associated macrophages [125]. Similarly, Chakarov *et al*.[25] highlighted selective expression of *Il10* in interstitial lung macrophages, and differential expression in the LYVE1^hi^/MHCII^lo^ subpopulation. They did not comment upon the reciprocal pattern observed with *Tnf* and *Ilb*, both more highly-expressed in the LYVE1^lo^ macrophages. Both populations of interstitial lung macrophages (and all the samples from other tissues in this BioProject) expressed very high levels of all of the IEG transcripts in Cluster 41. Whereas macrophage-expressed transcripts such as *Adgre1* are readily detected in total tissue mRNA, and are CSF1R-dependent, inflammatory cytokines and IEG transcripts are not [16,61]. Accordingly, in each of these studies, the expression of IEG and inducible cytokines is most likely an artefact of tissue disaggregation. Consistent with that conclusion, the clear exception in which IEG were not detected is peritoneal macrophages, which are not subjected to the stress of enzymic digestion during isolation.

Interestingly, *Acod1*, which was massively-induced within 1 hour by LPS in mouse macrophages *in vitro* (see http://biogps.org) was only detected at low levels in a small subset of samples and not correlated with IEG or any other inflammatory activation markers. Induction of this gene has attributed functions in adaptive immunometabolism and accumulation of TCA cycle intermediates in activated macrophages [126]. The lack of detection in the isolated macrophages suggests either that induction is specific to recruited inflammatory macrophages or that inducible expression is purely an *in vitro* phenomenon. The *Acod1* expression pattern was correlated only with *Il23a* (encoding a subunit of the cytokine IL23) at the stringency used here (*r* ≥ 0.75).

### Contamination of macrophage populations with other cell types

**Table 3** and **Figure 4B** highlight other clusters that were tissue-specific and contained markers and transcription factors associated with organ/tissue-specific differentiation, with corresponding enrichment for GO terms associated with specific tissues (**Table S2**). There are three ways in which mRNA from purified macrophage/DC populations may be contaminated with mRNA from unrelated cells. The most straightforward is poor separation of macrophages from unrelated contaminating cells by FACS for purely technical reasons. A second source derives from active phagocytosis by macrophages of senescent cells, where RNA from the engulfed cell may be detected. Finally, there is a phenomenon that arises from the extensive ramification of macrophages and their intimate interactions with other cells. Gray *et al*. [127] found that cells purified from lymph nodes with the surface marker CD169 were in fact lymphocytes coated with blebs of macrophage membrane and cytoplasm. Similarly, Lynch *et al*. [128] found that all methods to isolate Kupffer cells (KC) for flow cytometry produced significant contamination with CD31^+^ endothelium tightly adhered to remnants of KC membrane.

**Table 3.**
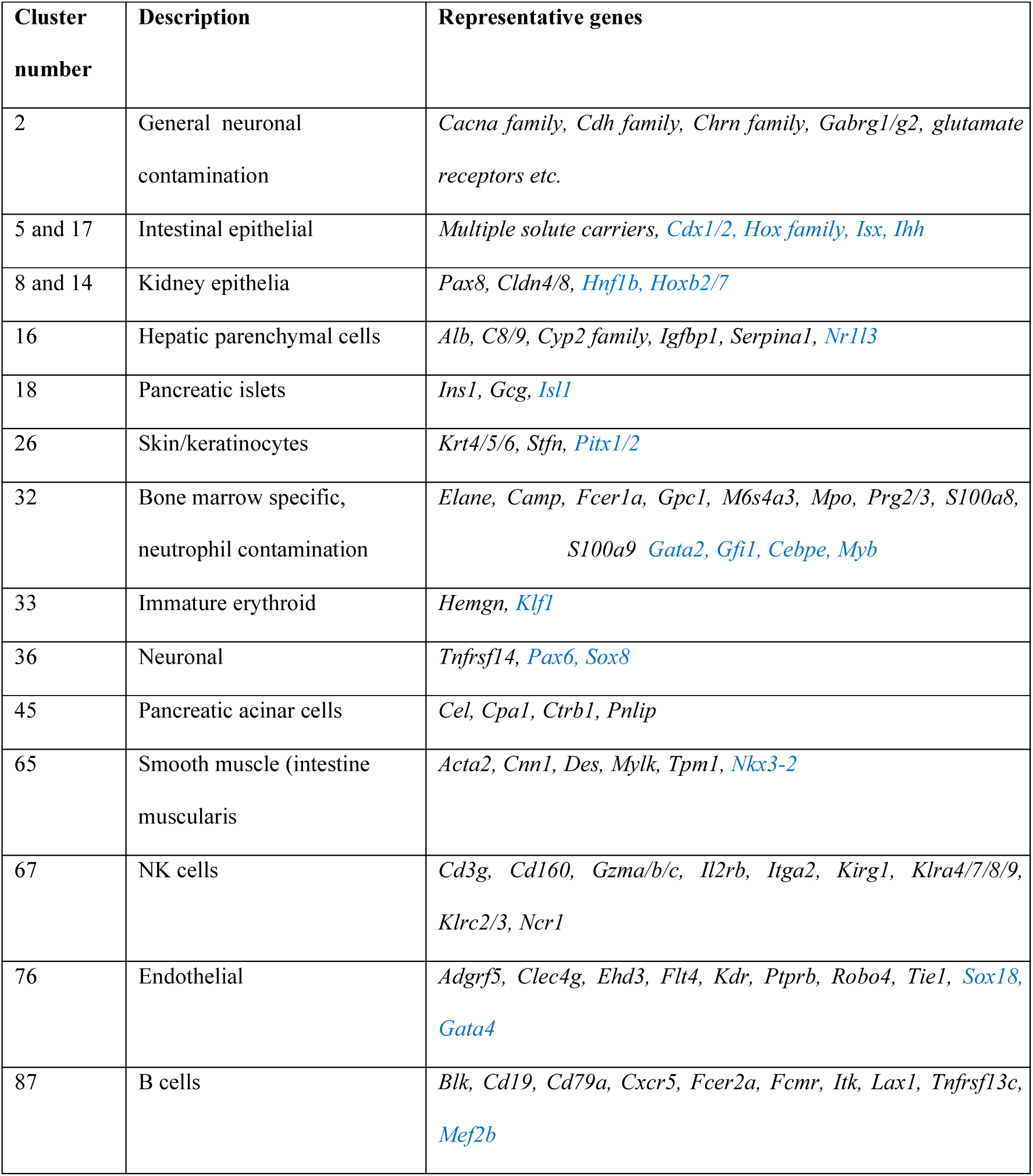
Major contaminant clusters. Genes in blue are transcription factors

Cluster 2 appears to be generic “rubbish” cluster, containing transcripts detected at relatively low levels only in specific BioProjects and unrelated to tissue of origin. Other clusters were driven by a single RNA-seq result from within one BioProject. These clusters most likely represent technical noise as well as contamination.

Consistent with the proposal from Lynch *et al*. [128] three endothelial-associated transcripts, *Cdh5, Pecam1* and *Stab 2*, were contained with the KC-enriched cluster (Cluster 12) and apparently increased in expression during KC differentiation. However, other endothelial transcripts were. Bonnardel *et al*. [129] generated RNA-seq data from purified liver sinusoidal endothelial cells (EC). We examined the profiles of the most highly-expressed EC genes in the macrophage dataset. Many of them were detectable in isolated KC but at much lower levels than *Cdh5, Pecam1* and *Stab2*. They contributed to a separate liver-specific endothelial cluster (Cluster 76). So, whilst there is evidence that EC contaminate KC preparations reflecting the close apposition in the sinusoids, *Chd5, Pecam1* and *Stab2* are likely also genuine KC-expressed transcripts.

The detection of mature red cell transcripts encoding haemoglobin (*Hba, Hbb*), which are quite abundant in many macrophage populations, most likely reflects ongoing erythrophagocytosis. Macrophages isolated from the intestinal lamina propria in one of the two large datasets from small intestine [33] were heavily contaminated with markers of intestinal epithelium (Clusters 5/17). This might be a separation artefact but could also reflect an active role of macrophages in clearance of senescent epithelial cells [80]. Cluster 18 and Cluster 45 were restricted to samples from a study of pancreatic islet and peri-islet macrophage populations [130]. The authors noted the expression of insulin (*Ins1*) mRNA in their islet macrophage populations and attributed it to an intimate interaction with β-cells. Contamination or β-cell-macrophage fusion was said to be excluded on the basis that β-cell markers such as *Pdx1* were not detected. However, many other islet-associated transcripts were abundant and formed part of Cluster 18, notably transcription factors *Isl1, Foxa2, Nkx6*.*1, Nkx2*.*2* and other abundant islet-specific transcripts, *Inhba, Chga/b, Iapp, Gipr* and *Gcg*. Similarly, Cluster 45 was relatively enriched in the peri-islet macrophages and contains transcripts encoding many pancreatic enzymes. Cluster 65 includes *Acta2* and other smooth muscle markers which selectively contaminated macrophages isolated from the intestinal muscularis [33].

The bone marrow contains several populations of macrophages [29] including those associated with hematopoietic islands expressing CD169 (*Siglec1* gene) and VCAM1. One of the datasets included in the present study profiled the transcriptome of macrophages associated with erythroblastic islands, based upon isolation using an *Epor-*EGFP reporter gene [131]. A second bone marrow dataset separated macrophages based upon their engagement in phagocytosis of blood-borne material [132]. The putative erythroblastic island macrophages did not actually express increased *Epor* mRNA (although *Epor* was detected in other macrophage populations as reported recently [68] and fell within the Cluster 22). However, in the isolated bone marrow macrophages, *Siglec1* was correlated with high levels of both immature neutrophil (Cluster 32) and erythroid-associated (Cluster 33) mRNAs. The separation of these two clusters implies that the contamination occurs in distinct macrophage populations, enriched selectively in each preparation and perhaps derived from separate hematopoietic islands [29]. Cluster 32 also contains the myeloid progenitor transcription factor *Myb* and the GM-progenitor marker *Ms4a3*. Given the extensive ramification of marrow macrophages and their intimate interactions with progenitors [29], this contamination likely reflects the same isolation artefact reported in lymph node [127], namely haemopoietic progenitor cells cloaked in macrophage clothing.

There are separate clusters including B cell and NK cell-specific markers. The B cell cluster, Cluster 87, shows highest average expression in intestine, bone marrow, lung and spleen and likely reflects close association between macrophages and B cells in lamina propria and germinal centres [33]. The cluster containing NK cell markers, Cluster 67, had highest average expression in one of the DC preparations. Those DC came from a study that proposed a further subdivision of cDC2 based upon expression of transcription factors T-bet (*Tbx21*) and RORγT [133] and separated cDC2 based upon expression of a *Tbx21* reporter allele. *Tbx21* was detected in all purified splenic cDC preparations presented on http://biogps.org, but at much lower levels than in NK cells. NK cells also express *Itgax*, used in purification of the cDC. Accordingly, it seems likely that apparent *Tbx21* expression in DC is due to NK cell contamination.

### Clustering of transcription factor (TF) expression

Most of the co-regulated clusters identified above contain genes encoding transcriptional regulators that are known to be essential for tissue-specific adaptation. These represent only a small subset of the transcriptional factors detected in MPS cells. The *r* value of 0.75 was chosen empirically for the analysis of the whole dataset to maximise the number of genes included while minimizing the number of edges between them (**Figure S1**) and aimed at assessing the predictive value of markers including those shown in **Figure 2**. To test the effect of reducing the stringency we focused on annotated transcription factors [134] to reduce the complexity and remove noise. 1103 transcriptional regulators were detected above the 10 TPM threshold in at least one macrophage population. The sample-to-sample matrix (**Figure 6**) shows that populations sourced from different tissues could be distinguished based upon TF expression alone. It also shows that TF expression in the DC populations was similar to that in macrophage cells. We generated GCN at three different Pearson correlation coefficient thresholds, 0.5, 0.6 and 0.7. The results are provided in **Table S3**. As the cut-off was reduced, more TF transcripts were included in the network. At the highest stringency *r* value ≥ 0.7, the largest cluster includes *Spi1* alongside many of the transcription factors identified in the largest generic MPS co-expression clusters above (Clusters 1, 3 and 4). We conclude that the basic shared identity of MPS cells involves coordinated expression of around 100-150 transcription factors. Even at the lowest *r* value (≥ 0.5), transcription factor genes identified as specific to particular tissue-specific MPS populations made few additional connections, indicating that local adaptation is dependent on highly-correlated and regulated expression of a small cohort of TF. Nevertheless, associations that become evident at lower *r* value may identify combinatorial interactions in particular cell populations; *Mycl*, associated with DC fitness ([135] was weakly-correlated with *Irf8* and *Zbtb46*; *Cebpb* with *Nfil3* and interferon-related transcription factors (*Batf2, Irf1/7/9, Stat1/2*) were connected at the threshold of 0.5 (**Table S3**).

**Figure 6.**
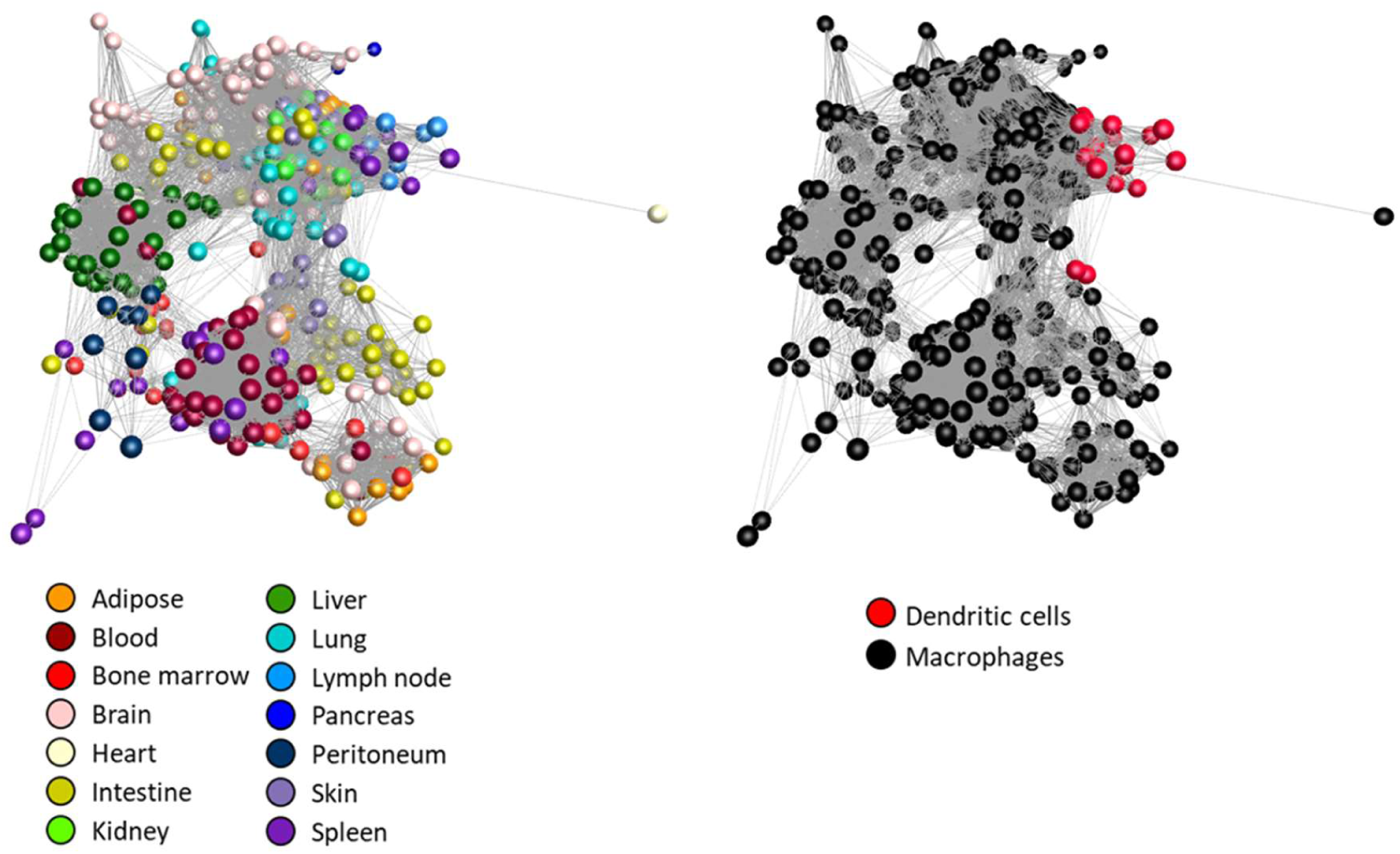
Network analysis of transcription factor gene expression in MPS cell populations. The sample-to-sample network was generated by Graphia analysis, at *r* ≥ 0.66, which included all 466 samples. Nodes representing samples are coloured by source tissue (left) and cell type (right).

### Expression of solute carriers and metabolism genes in macrophage populations

The burgeoning field of immunometabolism has focused on regulation of intermediary metabolism in recruited monocytes and macrophages in various states of activation or polarization [126]. Amongst emerging concepts is the view that M1 polarization (classical activation) is associated with aerobic glycolysis and mitochondrial dysfunction, whereas M2 polarization requires an active tricarboxylic acid cycle [126]. Cluster 7 contains mitochondria-associated transcripts and transcripts encoding ribosomal subunits, with variable expression across all samples, even from the same tissue, indicating the resident tissue macrophages vary in their dependence upon mitochondrial oxidative phosphorylation irrespective of surface markers or differentiation state.

In many cases metabolic pathways are regulated at the level of solute transport [126]. There were > 400 members of the large solute carrier (SLC) family expressed in mononuclear phagocytes above the 10 TPM threshold. Some were more highly expressed in intestine and kidney epithelial cells and clustered with tissue-specific epithelial markers. However, many contributed to macrophage-enriched expression clusters. One such gene, *Slco2b1*, which encodes an organic anion transporter of unknown function, has been proposed as a marker gene to distinguish macrophages from DC subsets and the promoter was used in an inducible macrophage depletion strategy [25]. The larger dataset does not support this dichotomy. *Slco2b1* is part of Cluster 4, enriched in microglia and absent from multiple other macrophage populations as well as both cDC subsets.

Macrophages depend to varying degrees upon glutamine, glucose and fatty acids as fuels [136] and glutamine is an important immune regulator [137]. 14 different solute carriers from 4 families have been shown to transport glutamine [138]. Of the genes encoding these carriers, *Slc38a1* was widely expressed in MPS cells and did not fall within a cluster, whereas *Slc7a5, Slc7a7, Slc7a8* and *Slc38a7* were part of distinct macrophage-enriched clusters. Consistent with the importance of glutamine as a fuel for MPS cells, transcripts encoding enzymes of glutamine metabolism (*Gls, Glud1, Glul, Slc25a11*) were also highly-expressed and part of Clusters 1 and 3. By contrast, resident MPS cells apparently have limited expression of glucose transporters. *Slc2a1* (encoding glucose transporter GLUT1) was low, highly variable and idiosyncratic amongst tissues. A myeloid-specific conditional knockout of *Slc2a1* confirmed that GLUT1 is the major glucose transporter in macrophages but the loss of glucose as a fuel had remarkably little impact on macrophage function [139]. The expression of *Slc2a1* in cells isolated from tissues is difficult to interpret since the transporter is induced by hypoxia [140], which might arise during isolation, and *Slc2a1* was barely detectable in peritoneal macrophages, which are less likely to undergo stress during isolation. In the absence of *Slc2a1*, macrophages increase oxidation of fatty acids [139]. The *Slc27a1* gene, encoding the fatty acid transporter FATP1 which also contributes to functional regulation in macrophages [141, 142], was widely-expressed in tissue macrophages and, with carnitine acyl transferase genes (*Crat, Crot*), formed part of Cluster 1. *Slc2a5* (found in Cluster 4) encodes a fructose-specific transporter [143] and was expressed primarily in microglia. *Slc2a6* is a lysosome-associated glucose transporter that was recently knocked out in the mouse genome [144]. It also has a novel expression profile being highest in monocytes and cDC2.

One of the best known functional solute carriers in macrophages is natural resistance associated membrane protein 1 (NRAMP1; *Slc11a1* gene), which is associated with genetic resistance to intracellular pathogens. SLC11A1/NRAMP1 is expressed in lysosomes and contributes to pathogen resistance by restricting available iron [145]. The role in iron metabolism is reflected by its presence in Cluster 12, alongside *Slc40a1*, encoding ferriportin, the macrophage-enriched iron exporter [146]. One other prominent class of solute carriers highly expressed in macrophages (*Slc30a6, Slc30a7, Slc30a9; Slc39a3, Slc39a7, Slc39a9* in Cluster 1; *Slc39a12* in Cluster 4) is involved in transport of zinc, which is a component of antimicrobial defense [147, 148]. Two further zinc transporters, *Slc39a2* and *Slc39a11*, were enriched in lung macrophages (Cluster 10). This lung macrophage-enriched cluster also contains *Slc52a3*, encoding a riboflavin transporter, *Slc6a4* (sodium and chloride dependent sodium symporter) and 2 members of the Slc9 family of sodium-hydrogen exchange (NHE) transporters (*Slc9a4* and *Slc9a7*) which are more traditionally associated with epithelial function [149].

### Validation of co-expression clustering with an independent kidney dataset

The abundant macrophage populations of the kidney were first described in detail using F4/80 as a marker *in situ* [150]. There has been considerable debate about the relationships between resident macrophages, monocyte-derived macrophages and cDC subsets in the kidney [151]. The main cluster analysis did not reveal a separate kidney resident macrophage-enriched profile. The kidney dataset in the analysis included F4/80^+^, CD64^+^ macrophages isolated from control and ischemic kidneys, further subdivided based upon expression of CD11B and CD11C [152]. Salei *et al*. [111] recently produced RNA-seq data for isolated populations of resident macrophages, monocyte-derived cells, cDC1 and cDC2 from kidney compared to similar populations from spleen. The primary data were not available for download by our automated pipeline through the ENA at the time we pooled and froze our dataset (February 2020). We therefore obtained the processed data directly from the authors and carried out network analysis using the 33 samples and 9,795 genes with normalized expression of at least 10 in at least one sample. The macrophages of the kidney are intimately associated with the capillaries [153] but *Lyve1* was not detectable in resident macrophages in this dataset or in [152]. Published IHC on mouse kidney reveals that LYVE1 is restricted to lymphatic vessels [154].

**Figure 7** illustrates the way in which the sample-to-sample matrix revealed relationships between the cell populations with increasing correlation coefficient threshold. Even at the lowest correlation cut-off, used in the main atlas (0.75), the splenic red pulp macrophages separated from all kidney and DC samples. As the cut-off was made more stringent, the cDC1 from both spleen and kidney separated, but the resident kidney macrophages, cDC2 and monocyte-derived macrophages remained closely connected until r ≥ 0.98 when the spleen cDC2 separated from the monocyte-derived macrophages and kidney cDC2. At r ≥ 0.99 the kidney cDC2 and monocyte-derived macrophages were still not separated indicating that the expression profiles of these cell types are very similar. Salei *et al*. [111] performed a principal components analysis based upon the 500 most variable transcripts and also identified the close relationship between cDC2 and monocyte-derived cells. Our analysis further emphasizes their conclusion that the main axis of difference is between spleen macrophages and all other cells. cDC1 from both tissues were more similar to each other than to the other cells but spleen cDC2 were only separated from kidney cDC2 and monocyte derived macrophages at the highest stringencies. We also performed a gene-to-gene analysis on these data. The profiles of kidney myeloid cells other than cDC1 were very similar and differed by only a small number of genes. Consistent with this conclusion, the two largest clusters in this analysis (see **Table S4**) were shared between all of the isolated populations and contain *Spi1* as well as many of the DC-enriched markers identified in the main analysis. However, *Ccr7* and many of the genes associated with it in the main dataset (Cluster 13, **Table 2**; e.g. *Spib, Stat4, Vsig10, Cd200, Itgb8*) were expressed at low levels in isolated kidney DC as in lung DC. Cluster 3 of the kidney analysis was specific to splenic red pulp macrophages and contains the known transcriptional regulators *Pparg, Spic* and *Nr1h3*. Transcripts in Cluster 4 were enriched in the resident kidney macrophages compared to both splenic macrophages and other kidney myeloid populations. Interestingly, the resident kidney macrophage cluster includes many genes that are also highly-expressed in microglia and depleted in the brain in *Csf1r* mutant mice and rats, including *Cx3cr1, C1qa/b/c, Csf3r, Ctss, Fcrls, Hexb, Laptm5, Tgfbr1, Tmem119* and *Trem2* [16, 61]. These were also detected in the isolated kidney macrophages in **Table S1**. Both microglia and resident F4/80^hi^ kidney macrophages are selectively lost in a mouse line with a mutation in a conserved enhancer of the *Csf1r* locus [16]. *Runx1*, which regulates the activity of the *Csf1r* enhancer [155] and has also been implicated in the establishment of microglial cells during development [156] was within this cluster. *Csf1r* mRNA was expressed at high levels in cells defined as cDC2 as well as monocyte-derived cells and resident macrophages. All cells expressing a *Csf1r*-EGFP reporter in the kidney were depleted by treatment with anti-CSF1R antibody[30]. This suggests that despite their expression of FLT3, renal cDC2 are CSF1R-dependent. Cluster 6 of the kidney analysis, including *Itgam*, was enriched in the selected CD11B^+^ populations from kidney but highly-expressed in all of the populations. This cluster includes all of the co-regulated immediate early genes identified in Cluster 41 in the extended MPS dataset above, suggesting that recent monocyte-derived cells may be more susceptible to activation during isolation.

**Figure 7.**
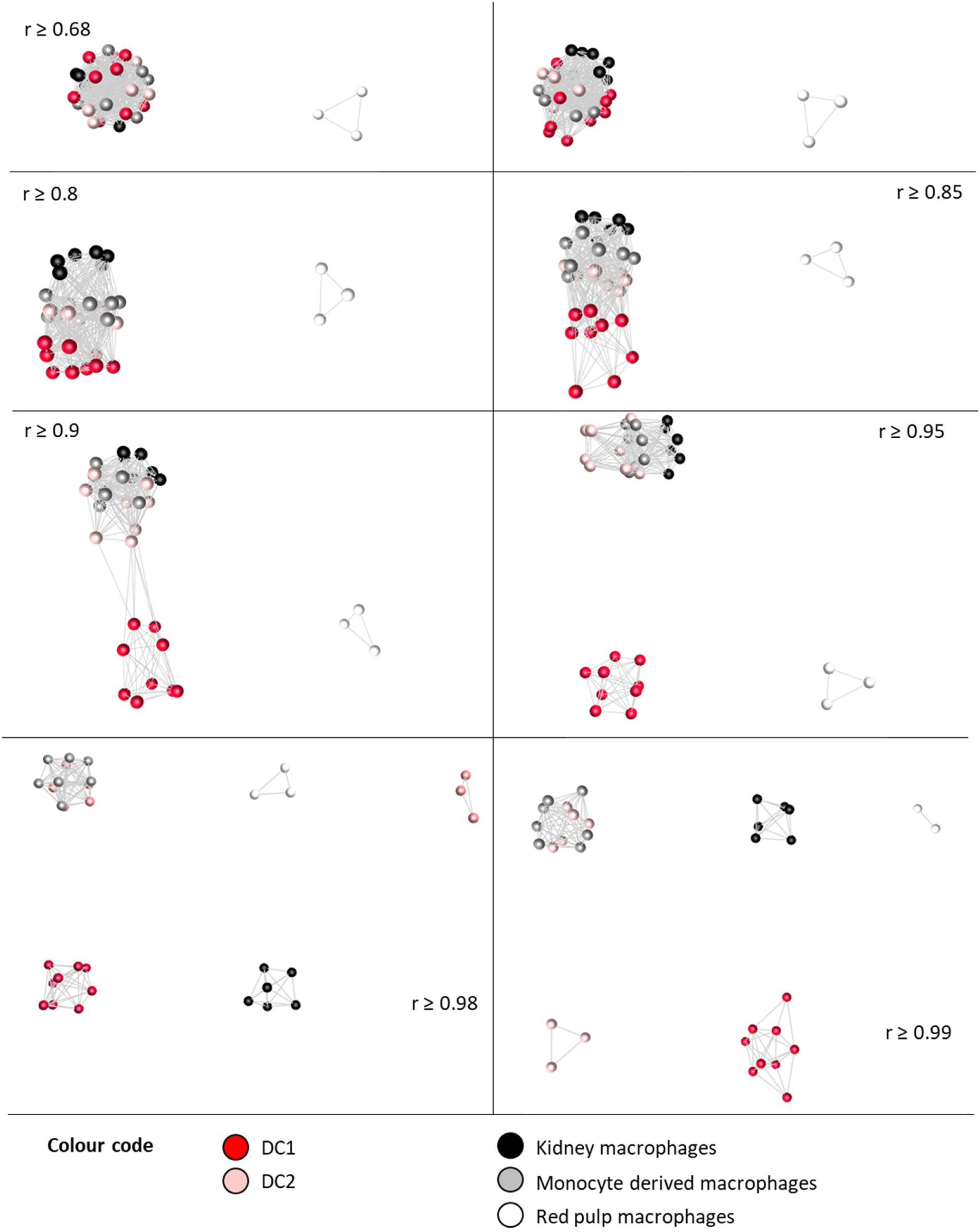
Network analysis of gene expression in macrophage and DC subpopulations from kidney. The sample-to-sample network was generated by Graphia analysis, at the indicated *r* values which all included all 33 samples up to *r ≥* 0.98. At *r ≥* 0.98 one red pulp macrophage sample was lost.

### The relationship between single cell and bulk RNA-seq data

The advent of scRNA-seq has been heralded as a revolution promising new approaches to classification of myeloid heterogeneity [35, 157, 158]. Single cell (sc) RNA-seq is intrinsically noisy, non-quantitative stochastic sampling of a subset of the most abundant mRNAs in individual cells ([159, 160]). Algorithms that support non-linear dimensional reduction (e.g. *t*-distributed stochastic neighbour embedding [*t*-SNE] or Uniform Manifold Approximation and Projection (UMAP)) [161] followed by some form of clustering are then used to join together groups of cells in which the members share detection of an arbitrarily-defined set of markers. There is an implicit assumption in this approach that defined cell types, with approximately identical transcriptomes, actually exist and that sampling noise can be overcome by pooling the transcripts detected in a sufficiently large number of cells to create meta-cells. Based upon scRNA-seq analysis of interstitial lung macrophages, Chakarov *et al*.[25] inferred the existence of a subpopulation that expressed LYVE1. They then generated bulk RNA-seq data from separated LYVE1^hi^ and LYVE1^lo^ subpopulations. Their data allow a critical comparison of the two approaches. For this purpose, the primary scRNA-seq data were downloaded, reanalysed and expressed as TPM using the Kallisto pseudoaligner. **Table S5** contains these reprocessed data, alongside the bulk RNA-seq data for the lung macrophage subpopulations, with the level of expression ranked based upon the bulk RNA-seq data for the purified LYVE1^hi^ interstitial macrophages.

Consistent with Zipf’s law, the power-law distribution of transcript abundance [162, 163], the top 200 expressed transcripts in the bulk RNA-seq data contribute around 50% of the total detected transcripts in the scRNA-seq data and it is clear that these are the only transcripts detected reliably (**Table S5**). The abundant transcripts from bulk RNA-seq that are also detected in scRNA-seq samples include many cell type-specific surface markers which explains the ability to use scRNA-seq to discover such markers. These abundant transcripts also include IEG such as *Dusp1, Egr1, Fos, Ier2* and *Junb*, indicative of the activation that occurs during isolation as discussed above. The inducible cytokines including *Ccl2, Tnf, Il1b, Il6* and *Il10* were each detected in a subset of the cells. Of the most highly-expressed transcripts only a very small subset (e.g. *Actb, Apoe, B2m, Ccl6, Cd74, Ctsb, Fth1, Ftl1, Lyz2*) had non-zero values in all cells. The average expression of the top 500 transcripts in the single cells was similar to the bulk RNA-seq but the detected expression level varied over 4 orders of magnitude among individual cells. *Fcgr1* and *Mertk* mRNAs, encoding markers used to purify the interstitial macrophages for scRNA-seq, were actually detected in only a small subset of the cells and were not correlated with each other. Both this study and a subsequent study [50] state that *Mrc1* and *Lyve1* expression is shared by overlapping populations of lung interstitial macrophages. That conclusion is not supported by the data. The separation of these two markers was evident from the separate study of lung interstitial macrophage populations [164] included in our analysis and has been discussed above. Even in the bulk RNA-seq data from lung interstitial macrophages, the expression of *Mrc1* was only marginally-enriched in purified LYVE1^hi^ cells relative to LYVE1^lo^ cells (**Table S1**). Consistent with that conclusion, in the scRNA-seq data the two are not strictly correlated with each other, with *Mrc1* being detected in many more cells than *Lyve1* (**Figure 8A**) despite similar absolute levels of expression in the total RNA-seq data.

**Figure 8.**
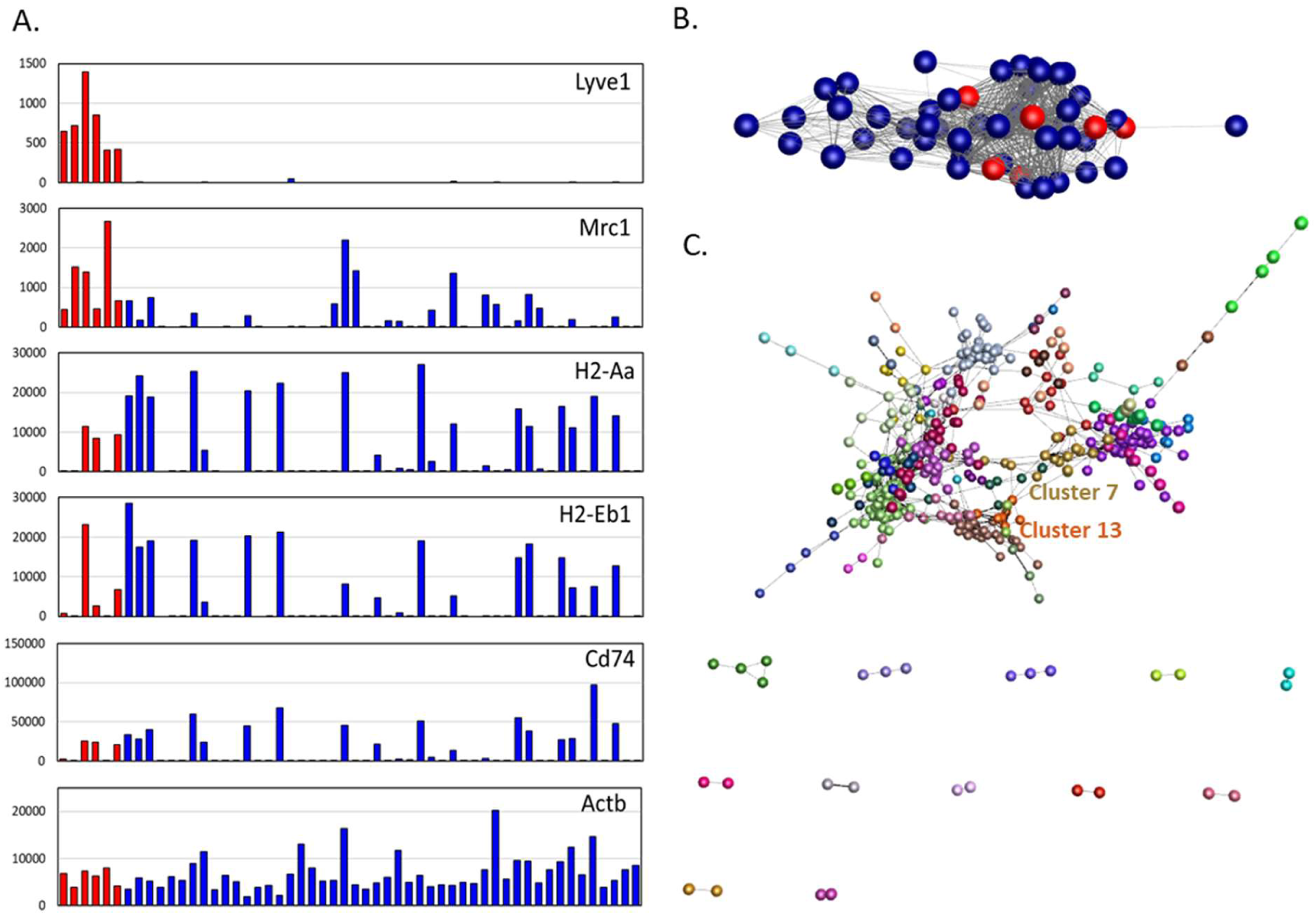
Network analysis of single cell RNA-seq data. **A**. Expression profiles in single cells for selected genes. Only the first six cells expressed *Lyve1* (coloured red). **B**. The sample-to-sample network was generated by Graphia analysis, at *r* ≥ 0.53, which included all 54 samples. Nodes represent samples; red nodes show the samples with high expression of *Lyve1*. **C**. Gene-to-gene network (*r* ≥ 0.5), clustered at MCL inflation value of 1.7. Cluster lists and expression profiles are available in **Table** S6.

To identify whether any robust correlations actually exist in the scRNA-seq data, the top 500 expressed transcripts in the scRNA-seq samples were used for network analysis. The sample-to-sample network (*r*≥0.53) is shown in **Figure 8B** and the gene-to-gene network (*r*≥0.5) in **Figure 8C**. The cluster list and average expression profiles are provided in **Table S6**. One clear-cut finding is the co-expression of genes involved in APC activity (*H2-Aa, H2-Ab1, H2-Eb1, Cd74* and *Ctss*; Cluster 13 of the scRNA-seq analysis), which were effectively present or absent in individual cells. Chakarov *et al*. [25] defined two subpopulations as LYVE1^hi^/MHCII^lo^ and LYVE1^lo^/MHCII^hi^ but only six of the scRNA-seq samples expressed *Lyve1* and half of those also expressed MHC II genes (**Figure 8A**). This is consistent with the lack of any inverse correlation between *Lyve1* and *Cd74* in **Figure 2B**. Even at this low *r* value known highly-expressed markers segregated from each other. *Lyve1* forms a cluster with *Mgl2, Cd209 and Cd302* (Cluster 7 in the scRNA-seq analysis; **Figure 8C**). *Adgre1* is in a co-expression cluster that includes *Lyz2* and *Msr1* (Cluster 4 of the scRNA-seq analysis), *Csf1r* is co-expressed with *Mrc1* and *Cd163* (Cluster 2) and *Lgals3* with *Retnla* and *Fcrls* (Cluster 1). The co-regulation of MHC-related genes, and genes located in the same chromosomal region (e.g. *C1qa, C1qb, C1qc*; Cluster25 of the scRNA-seq analysis) as well as the relatively uniform detection of genes such as *Actb* (**Figure 8A**) suggest that a significant proportion of the all-or-nothing differences in expression between cells in the scRNA-seq data is real.

### Implications of transcriptional network analysis for the utility of surface markers

As discussed in the previous section, scRNA-seq provides an ambiguous view of cellular heterogeneity. Non-linear dimensionality reduction algorithms can hide or emphasise diversity by grouping cells in which an overlapping set of markers is detected. The number of populations defined depends upon the parameters applied and different approaches do not always give the same answers [161]. In the lung interstitial population we reanalysed, the detected expression of transcripts encoding plasma membrane proteins was essentially all-or-nothing. That conclusion may be considered a reflection of the limitations of the technology, but it is actually supported by other evidence. Tan and Krasnow [165] defined subpopulations of interstitial lung macrophages based upon expression of F4/80, Mac-2 (*Lgals3*) and MHCII, and tracked the changes in their relative abundance during development. Interestingly, they did not detect LYVE1 on adult lung interstitial macrophages by IHC. Consistent with their data, in the scRNA-seq data most lung interstitial macrophages expressed high levels of either *Adgre1* or *Lgals3*, but some expressed both or neither. Protein expression at a single cell level clearly does not vary to the same extent as mRNA since proteins have different rates of turnover and decay [166]. Markers such as F4/80 and CD11C, and transgenes based upon macrophage-enriched promoters such as *Csf1r* and *Cx3cr1* do appear to label the large majority of MPS cells in most tissues. The disconnect between scRNA-seq and cell surface markers may partly reflect the nature of transcription. At the single cell level transcription occurs in pulses interspersed by periods of inactivity and mRNA decay, which can manifest as random monoallelic transcript expression [167]. If gene expression is genuinely probabilistic at the level of individual loci [166] the assumption of transcriptomic homogeneity in definable cell types upon which scRNA-seq analysis is based is clearly invalid. The number of macrophage subpopulations that can be defined in any scRNA-seq dataset becomes a matter of choice and model. As an extreme example, one recent scRNA-seq study identified 25 distinct myeloid cell differentiation “states” in a mouse lung cancer model [168].

The RNA-seq data included as representative of cDC subsets [133] was from cells purified using CD64 as a marker to exclude macrophages. Despite this choice, an unbiased assessment of the sample-to-sample matrix in **Figure 3B** (based on all genes) and **Figure 6** (based on transcription factor genes) would class all of these DC as part of the same family as macrophages. The use of CD64 as a definitive marker distinguishing macrophages from DC was criticized when it was proposed [42] and it remains untenable. It is actually a curious choice as a marker to define a cell as a macrophage since the protein FCGR1 (CD64) has been implicated functionally in APC activity [169]. From our analysis, the clear separation of DC from all other members of the MPS based upon APC function, surface markers, transcription factors or ontogeny [20] remains problematic. The one criterion that remains is location. We suggest that the only defensible definition of a cDC is a mononuclear phagocyte that is adapted specifically to the T cell areas of secondary lymphoid organs, responding to a specific growth factor (FLT3L) and the chemokine receptor CCR7. The kidney data [111] suggest that there is also tissue-specific adaptation of “cDC2” which may remain more “macrophage-like”. In that sense cDC1 and cDC2 are no more unique than a peritoneal macrophage adapting to signals from retinoic acid via induction of *Gata6*.

### A critical view of markers of macrophage polarization states

The concept of M1/M2 polarization derives from analysis of classical and alternative activation of recruited monocytes by Th1 (γIFN) and Th2 (IL4/IL13) cytokines [74]. Previous meta-analysis indicated that proposed M2 markers defined by others [74] correlate poorly with each other in isolated inflammatory macrophages and are not conserved across species [28]. The M1/M2 concept was also challenged in a recent comparative analysis of *in vitro* and *in vivo* data on macrophage gene expression [170] which concluded that “valid *in vivo* M1/M2 surface markers remain to be discovered”. We would suggest that they do not exist. The pro-inflammatory cytokines that we identified as inducible in all the resident MPS cells during isolation would be considered indicators of M1-like activation. Aside from proposed M2 markers already mentioned that each have idiosyncratic expression (*Mrc1, Retnla, Igf1, Mgl2*), *Chil3* (aka Ym1) was highly-expressed in lung macrophages (Cluster 10 of the whole dataset analysis; **Table S2**), *Arg1* and *Alox15* were restricted to peritoneal macrophages (Cluster 21) and *Cd163* was part of a small cluster of 4 transcripts (Cluster 312). The cluster analysis indicates that the detection of M2 markers on resident tissue macrophages has little predictive value. Detection of CD206 cannot imply that the cells have been stimulated with IL4/IL13, nor that they share any functions with alternatively-activated recruited monocytes. Nevertheless, IL4/IL13/STAT6 signalling could contribute to resident MPS cell differentiation. The IL13 receptor (*Il13ra*) is part of the generic MPS Cluster 1 and *Il4ra* is also highly and widely expressed. IL4 administration to mice can drive resident tissue macrophage proliferation beyond levels controlled homeostatically by CSF1[171].

### How do transcriptional networks contribute to understanding macrophage heterogeneity *in situ*?

An important question that arises from the transcriptomic analysis of subpopulations of resident MPS cells is precisely where are they located and how do they relate to each other? A significant concern with analysis of the cells isolated by tissue digestion and analysed here is whether recovered cells are representative of the tissue populations. Analysis of the lung has suggested that interstitial macrophages are a minority of the lung macrophage population [164]. That conclusion is not compatible with our own studies visualising *Csf1r* reporter genes, where the stellate interstitial populations are at least as abundant as alveolar macrophages [48, 49]. The description of subpopulations is not often linked to precise location with the tissue. One exception is the apparent location of LYVE1^hi^ macrophages with capillaries in the lung [25]. On the other hand, it is unclear where the putative long-lived CD4^+^, TIM4^+^ population in the gut [34] is located. In broad overview, macrophages in every organ, detected with *Csf1r* reporter transgenes that are expressed in all myeloid cells including DC, have a remarkably regular and uniform distribution. The concept of a macrophage territory [5] or a niche [172, 173] has been proposed. But despite their apparent homogeneity in location and morphology, multicolour localisation of macrophage surface markers suggests that they are almost infinitely heterogeneous (reviewed in [23]). Most of the datasets analysed here suggest that monocytes and macrophages in each organ are a differentiation series. We take the view that macrophages in tissues have a defined half-life such that some cells survive by chance and continue to change their gene expression [5]. Each macrophage that occupies a new territory, either following infiltration as a monocyte or self-renewal by cell division, starts a life history that involves changes in gene expression and surface markers with time. In that view, MPS subpopulations are no more than arbitrary windows within a temporal profile of adaptation.

## Conclusion

The transcriptional network analysis confirms that using our unique approach to down-sizing and a common quantification pathway, the RNA-seq data from different laboratories can be merged to provide novel insights. The network analysis indicates the power of large datasets to detect sets of co-regulated transcripts that define metastable states of MPS adaptation and function. The merged dataset we have created provides a resource for the study of MPS biology that extends and complements resources such as ImmGen (http://www.immgen.org). It can be readily expanded to include any new RNA-seq data for comparative analysis. Clusters of transcripts that are robustly correlated give clear indications of shared functions and transcriptional regulation. However, our analysis also reveals two important artefacts in the study of isolated tissue macrophages, the clear evidence of inflammatory activation during isolation and the extensive contamination of isolated preparations with transcripts derived from other cell types.

A discussion review of MPS heterogeneity in 2010 [174] suggested that in order for the field of immunology to advance and communicate “all cells have to be called something”. This Linnaean view continues to drive efforts to classify MPS cells into subsets based upon markers. The analysis we have presented shows that surface markers are poorly associated with each other and have very limited predictive value. Aside from MHCII, there are no markers that can be correlated with predicted APC activity. Resident tissue MPS cells, including cells that are currently defined as DC, belong to a closely-related family of cells in which the transcriptomic similarities are much greater than the differences. The cumulative function of the population of MPS cells acting together within each tissue is likely to be more important to homeostasis and immunity than the individual heterogeneity.

## ACKNOWLEDGEMENTS

We would like to thank Dr Barbara Schraml, Ludwig-Maximilians-University of Munich, for providing access to data not yet available in the public domain. Mater Research Institute-University of Queensland is grateful for support from the Mater Foundation, Brisbane, Australia. The Translational Research Institute receives funding from the Australian Government.

---

**Figure S1.**
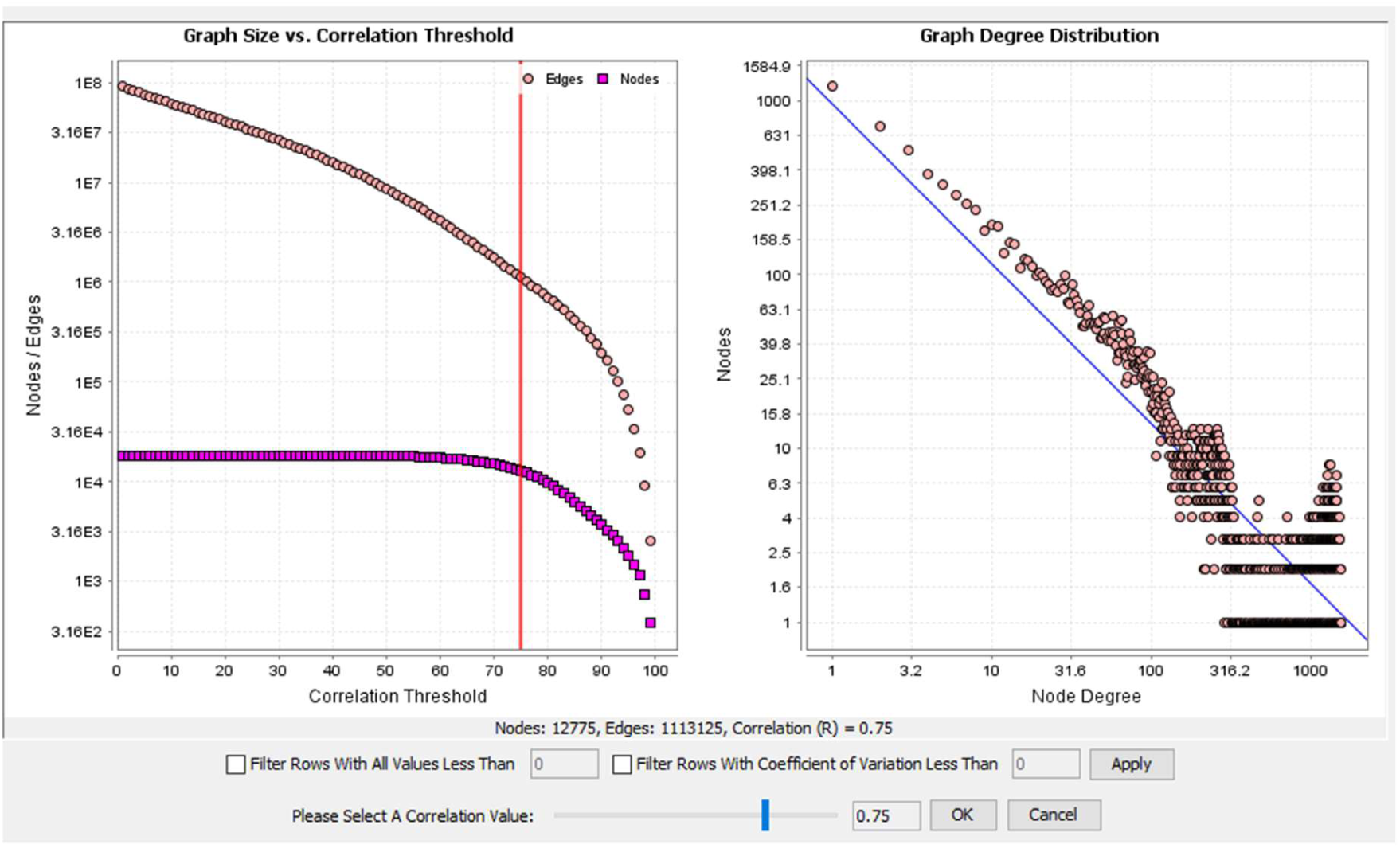
Graph size compared with correlation threshold for the analysis of the mouse macrophage dataset. The chosen correlation threshold of 0.75 resulted in inclusion of 12,775 nodes making 1,113,125 edges (correlations of ≥ 0.75) between them.

**Figure S2.**
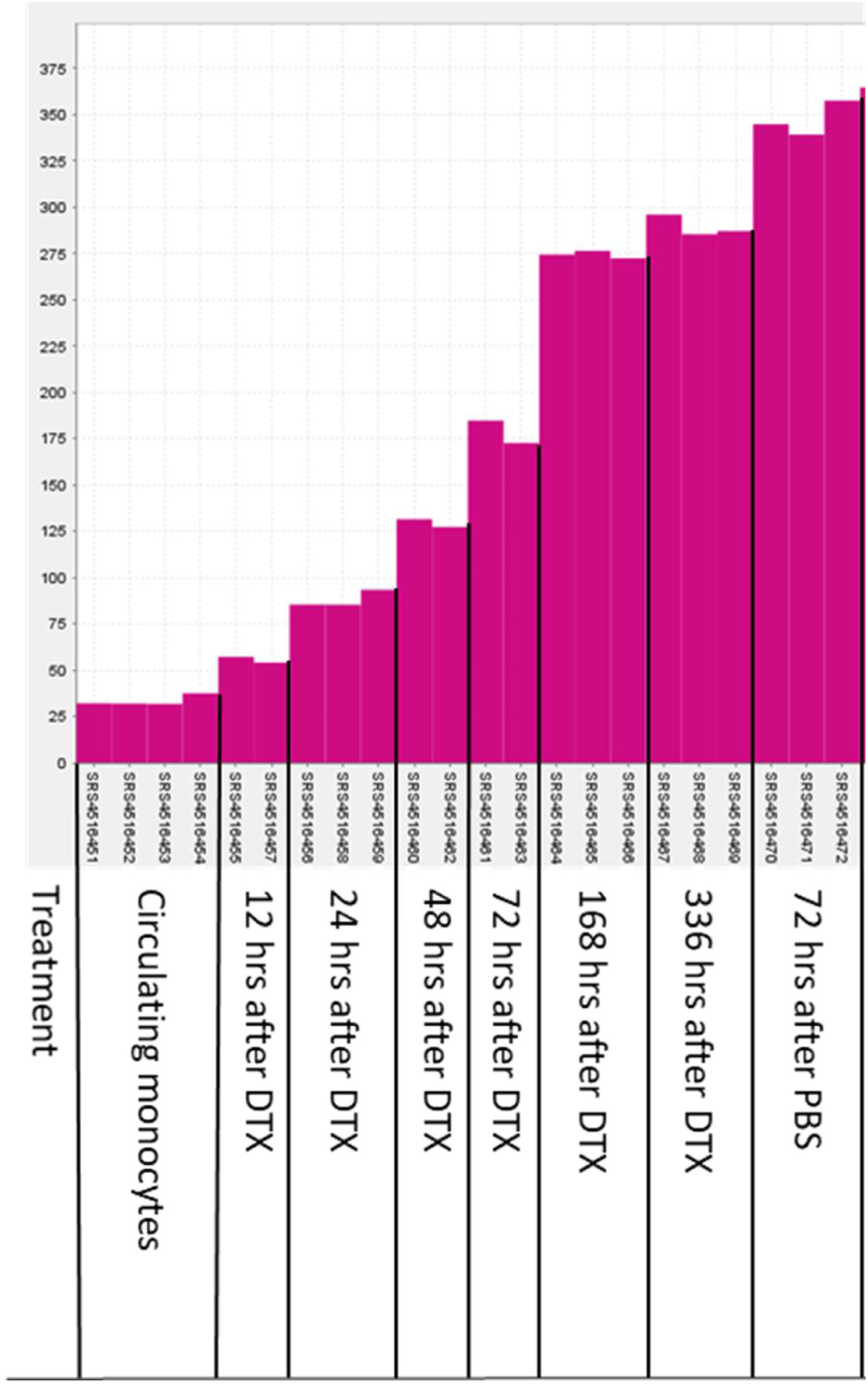
Average expression of genes in Cluster 12 during differentiation of monocytes to Kupffer cells. Data from BioProject PRJNA528435. *Clec4f*-cre Rosa26iDTX mice were treated with diptheria toxin (DTX) to remove mature Kupffer cells livers were harvested at indicated time points after DTX treatment. The experiment shows the repopulation of the liver with cells derived from monocytes. Control animals were treated with PBS and harvested at 72 hours.

**Figure S3.**
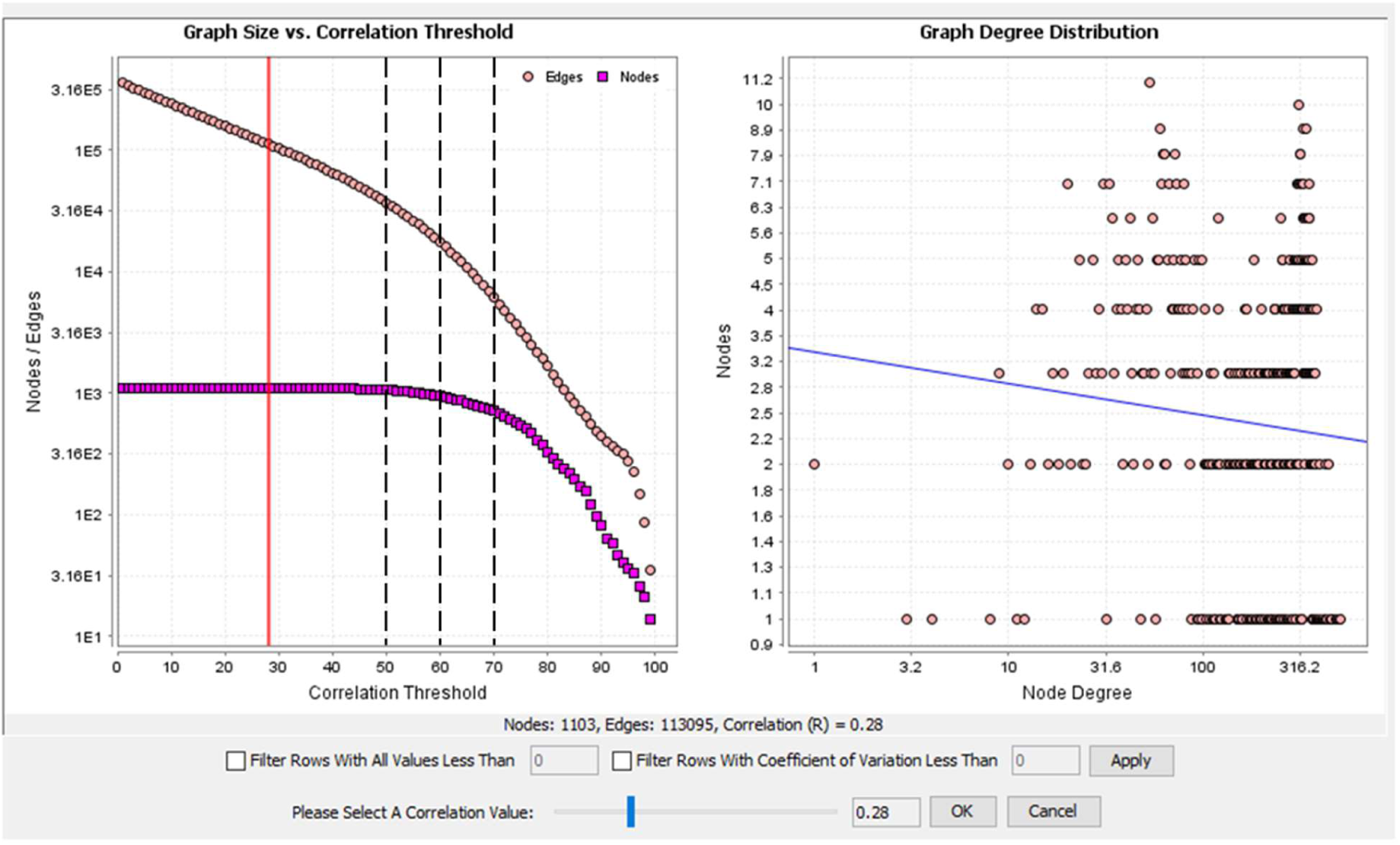
Graph size compared with correlation threshold for the analysis of the mouse macrophage transcription factor dataset. Red line shows the highest threshold to include all 1103 nodes (*r* ≥ 0.28). Black broken lines show the three correlation thresholds used in the analysis, 0.5 (1064 nodes), 0.6 (949 nodes) and 0.7 (714 nodes).

## References

1. van Furth, R., Cohn, Z. A., Hirsch, J. G., Humphrey, J. H., Spector, W. G., Langevoort, H. L. (1972) The mononuclear phagocyte system: a new classification of macrophages, monocytes, and their precursor cells. Bull World Health Organ 46, 845–52.

2. Amit, I., Winter, D. R., Jung, S. (2016) The role of the local environment and epigenetics in shaping macrophage identity and their effect on tissue homeostasis. Nat Immunol 17, 18–25.

3. Hoeffel, G. and Ginhoux, F. (2018) Fetal monocytes and the origins of tissue-resident macrophages. Cell Immunol 330, 5–15.

4. Hoeksema, M. A. and Glass, C. K. (2019) Nature and nurture of tissue-specific macrophage phenotypes. Atherosclerosis 281, 159–167.

5. Hume, D. A., Irvine, K. M., Pridans, C. (2018) The Mononuclear Phagocyte System: The Relationship between Monocytes and Macrophages. Trends Immunol. 40, 98–112

6. Chitu, V. and Stanley, E. R. (2017) Regulation of Embryonic and Postnatal Development by the CSF-1 Receptor. Curr Top Dev Biol 123, 229–275.

7. Hume, D. A., Caruso, M., Ferrari-Cestari, M., Summers, K. M., Pridans, C., Irvine, K. M. (2020) Phenotypic impacts of CSF1R deficiencies in humans and model organisms. J Leukoc Biol 107, 205–219.

8. Summers, K. M. and Hume, D. A. (2017) Identification of the macrophage-specific promoter signature in FANTOM5 mouse embryo developmental time course data. J Leukoc Biol 102, 1081–1092.

9. Ginhoux, F. and Guilliams, M. (2016) Tissue-Resident Macrophage Ontogeny and Homeostasis. Immunity 44, 439–449.

10. Liu, Z., Gu, Y., Chakarov, S., Bleriot, C., Kwok, I., Chen, X., Shin, A., Huang, W., Dress, R. J., Dutertre, C. A., Schlitzer, A., Chen, J., Ng, L. G., Wang, H., Liu, Z., Su, B., Ginhoux, F. (2019) Fate Mapping via Ms4a3-Expression History Traces Monocyte-Derived Cells. Cell 178, 1509–1525 e19.

11. Yona, S., Kim, K. W., Wolf, Y., Mildner, A., Varol, D., Breker, M., Strauss-Ayali, D., Viukov, S., Guilliams, M., Misharin, A., Hume, D. A., Perlman, H., Malissen, B., Zelzer, E., Jung, S. (2013) Fate mapping reveals origins and dynamics of monocytes and tissue macrophages under homeostasis. Immunity 38, 79–91.

12. Jenkins, S. J. and Hume, D. A. (2014) Homeostasis in the mononuclear phagocyte system. Trends Immunol 35, 358–67.

13. Bonnardel, J. and Guilliams, M. (2018) Developmental control of macrophage function. Curr Opin Immunol 50, 64–74.

14. Gomez Perdiguero, E., Klapproth, K., Schulz, C., Busch, K., Azzoni, E., Crozet, L., Garner, H., Trouillet, C., de Bruijn, M. F., Geissmann, F., Rodewald, H. R. (2015) Tissue-resident macrophages originate from yolk-sac-derived erythro-myeloid progenitors. Nature 518, 547–51.

15. Mass, E., Ballesteros, I., Farlik, M., Halbritter, F., Gunther, P., Crozet, L., Jacome-Galarza, C. E., Handler, K., Klughammer, J., Kobayashi, Y., Gomez-Perdiguero, E., Schultze, J. L., Beyer, M., Bock, C., Geissmann, F. (2016) Specification of tissue-resident macrophages during organogenesis. Science 353. aaf4238

16. Rojo, R., Raper, A., Ozdemir, D. D., Lefevre, L., Grabert, K., Wollscheid-Lengeling, E., Bradford, B., Caruso, M., Gazova, I., Sanchez, A., Lisowski, Z. M., Alves, J., Molina-Gonzalez, I., Davtyan, H., Lodge, R. J., Glover, J. D., Wallace, R., Munro, D. A. D., David, E., Amit, I., Miron, V. E., Priller, J., Jenkins, S. J., Hardingham, G. E., Blurton-Jones, M., Mabbott, N. A., Summers, K. M., Hohenstein, P., Hume, D. A., Pridans, C. (2019) Deletion of a Csf1r enhancer selectively impacts CSF1R expression and development of tissue macrophage populations. Nat Commun 10, 3215.

17. Henson, P. M. and Hume, D. A. (2006) Apoptotic cell removal in development and tissue homeostasis. Trends Immunol 27, 244–50.

18. Hume, D. A. (2008) Macrophages as APC and the dendritic cell myth. J Immunol 181, 5829–35.

19. Jakubzick, C. V., Randolph, G. J., Henson, P. M. (2017) Monocyte differentiation and antigen-presenting functions. Nat Rev Immunol 17, 349–362.

20. Guilliams, M., Ginhoux, F., Jakubzick, C., Naik, S. H., Onai, N., Schraml, B. U., Segura, E., Tussiwand, R., Yona, S. (2014) Dendritic cells, monocytes and macrophages: a unified nomenclature based on ontogeny. Nat Rev Immunol 14, 571–8.

21. Sichien, D., Lambrecht, B. N., Guilliams, M., Scott, C. L. (2017) Development of conventional dendritic cells: from common bone marrow progenitors to multiple subsets in peripheral tissues. Mucosal Immunol 10, 831–844.

22. Gordon, S. and Pluddemann, A. (2017) Tissue macrophages: heterogeneity and functions. BMC Biol 15, 53.

23. Hume, D. A. (2008) Differentiation and heterogeneity in the mononuclear phagocyte system. Mucosal Immunol 1, 432–41.

24. Hume, D. A. (2011) Applications of myeloid-specific promoters in transgenic mice support in vivo imaging and functional genomics but do not support the concept of distinct macrophage and dendritic cell lineages or roles in immunity. J Leukoc Biol 89, 525–38.

25. Chakarov, S., Lim, H. Y., Tan, L., Lim, S. Y., See, P., Lum, J., Zhang, X. M., Foo, S., Nakamizo, S., Duan, K., Kong, W. T., Gentek, R., Balachander, A., Carbajo, D., Bleriot, C., Malleret, B., Tam, J. K. C., Baig, S., Shabeer, M., Toh, S. E. S., Schlitzer, A., Larbi, A., Marichal, T., Malissen, B., Chen, J., Poidinger, M., Kabashima, K., Bajenoff, M., Ng, L. G., Angeli, V., Ginhoux, F. (2019) Two distinct interstitial macrophage populations coexist across tissues in specific subtissular niches. Science 363.eaau0964

26. Gautier, E. L., Shay, T., Miller, J., Greter, M., Jakubzick, C., Ivanov, S., Helft, J., Chow, A., Elpek, K. G., Gordonov, S., Mazloom, A. R., Ma’ayan, A., Chua, W. J., Hansen, T. H., Turley, S. J., Merad, M., Randolph, G. J., Immunological Genome, C. (2012) Gene-expression profiles and transcriptional regulatory pathways that underlie the identity and diversity of mouse tissue macrophages. Nat Immunol 13, 1118–28.

27. Miller, J. C., Brown, B. D., Shay, T., Gautier, E. L., Jojic, V., Cohain, A., Pandey, G., Leboeuf, M., Elpek, K. G., Helft, J., Hashimoto, D., Chow, A., Price, J., Greter, M., Bogunovic, M., Bellemare-Pelletier, A., Frenette, P. S., Randolph, G. J., Turley, S. J., Merad, M., Immunological Genome, C. (2012) Deciphering the transcriptional network of the dendritic cell lineage. Nat Immunol 13, 888–99.

28. Hume, D. A. (2015) The Many Alternative Faces of Macrophage Activation. Front Immunol 6, 370.

29. Kaur, S., Raggatt, L. J., Batoon, L., Hume, D. A., Levesque, J. P., Pettit, A. R. (2017) Role of bone marrow macrophages in controlling homeostasis and repair in bone and bone marrow niches. Semin Cell Dev Biol 61, 12–21.

30. MacDonald, K. P., Palmer, J. S., Cronau, S., Seppanen, E., Olver, S., Raffelt, N. C., Kuns, R., Pettit, A. R., Clouston, A., Wainwright, B., Branstetter, D., Smith, J., Paxton, R. J., Cerretti, D. P., Bonham, L., Hill, G. R., Hume, D. A. (2010) An antibody against the colony-stimulating factor 1 receptor depletes the resident subset of monocytes and tissue- and tumor-associated macrophages but does not inhibit inflammation. Blood 116, 3955–63.

31. Geissmann, F., Manz, M. G., Jung, S., Sieweke, M. H., Merad, M., Ley, K. (2010) Development of monocytes, macrophages, and dendritic cells. Science 327, 656–61.

32. Sakai, M., Troutman, T. D., Seidman, J. S., Ouyang, Z., Spann, N. J., Abe, Y., Ego, K. M., Bruni, C. M., Deng, Z., Schlachetzki, J. C. M., Nott, A., Bennett, H., Chang, J., Vu, B. T., Pasillas, M. P., Link, V. M., Texari, L., Heinz, S., Thompson, B. M., McDonald, J. G., Geissmann, F., Glass, C. K. (2019) Liver-Derived Signals Sequentially Reprogram Myeloid Enhancers to Initiate and Maintain Kupffer Cell Identity. Immunity 51, 655–670 e8.

33. De Schepper, S., Verheijden, S., Aguilera-Lizarraga, J., Viola, M. F., Boesmans, W., Stakenborg, N., Voytyuk, I., Schmidt, I., Boeckx, B., Dierckx de Casterle, I., Baekelandt, V., Gonzalez Dominguez, E., Mack, M., Depoortere, I., De Strooper, B., Sprangers, B., Himmelreich, U., Soenen, S., Guilliams, M., Vanden Berghe, P., Jones, E., Lambrechts, D., Boeckxstaens, G. (2019) Self-Maintaining Gut Macrophages Are Essential for Intestinal Homeostasis. Cell 175, 400–415

34. Shaw, T. N., Houston, S. A., Wemyss, K., Bridgeman, H. M., Barbera, T. A., Zangerle-Murray, T., Strangward, P., Ridley, A. J. L., Wang, P., Tamoutounour, S., Allen, J. E., Konkel, J. E., Grainger, J. R. (2018) Tissue-resident macrophages in the intestine are long lived and defined by Tim-4 and CD4 expression. J Exp Med 215, 1507–1518.

35. Bassler, K., Schulte-Schrepping, J., Warnat-Herresthal, S., Aschenbrenner, A. C., Schultze, J. L. (2019) The Myeloid Cell Compartment-Cell by Cell. Annu Rev Immunol 37, 269–293.

36. Hume, D. A. and Freeman, T. C. (2014) Transcriptomic analysis of mononuclear phagocyte differentiation and activation. Immunol Rev 262, 74–84.

37. Hume, D. A., Summers, K. M., Raza, S., Baillie, J. K., Freeman, T. C. (2010) Functional clustering and lineage markers: insights into cellular differentiation and gene function from large-scale microarray studies of purified primary cell populations. Genomics 95, 328–38.

38. Joshi, A., Pooley, C., Freeman, T. C., Lennartsson, A., Babina, M., Schmidl, C., Geijtenbeek, T., Consortium, F., Michoel, T., Severin, J., Itoh, M., Lassmann, T., Kawaji, H., Hayashizaki, Y., Carninci, P., Forrest, A. R., Rehli, M., Hume, D. A. (2015) Technical Advance: Transcription factor, promoter, and enhancer utilization in human myeloid cells. J Leukoc Biol 97, 985–995.

39. Mabbott, N. A., Baillie, J. K., Brown, H., Freeman, T. C., Hume, D. A. (2013) An expression atlas of human primary cells: inference of gene function from coexpression networks. BMC Genomics 14, 632.

40. Mabbott, N. A., Kenneth Baillie, J., Hume, D. A., Freeman, T. C. (2010) Meta-analysis of lineage-specific gene expression signatures in mouse leukocyte populations. Immunobiology 215, 724–36.

41. Doig, T. N., Hume, D. A., Theocharidis, T., Goodlad, J. R., Gregory, C. D., Freeman, T. C. (2013) Coexpression analysis of large cancer datasets provides insight into the cellular phenotypes of the tumour microenvironment. BMC Genomics 14, 469.

42. Hume, D. A., Mabbott, N., Raza, S., Freeman, T. C. (2013) Can DCs be distinguished from macrophages by molecular signatures? Nat Immunol 14, 187–9.

43. Bush, S. J., Freem, L., MacCallum, A. J., O’Dell, J., Wu, C., Afrasiabi, C., Psifidi, A., Stevens, M. P., Smith, J., Summers, K. M., Hume, D. A. (2018) Combination of novel and public RNA-seq datasets to generate an mRNA expression atlas for the domestic chicken. BMC Genomics 19, 594.

44. Summers, K., Bush, S., Wu, C., Su, A., Muriuki, C., Clark, E., Finlayson, H., Eory, L., Waddell, L., Talbot, R., Archibald, A., Hume, D. (2020) Functional annotation of the transcriptome of the pig, sus scrofa, based upon network analysis of an RNAseq transcriptional atlas. Frontiers Genetics. 10, 1355

45. Gal-Oz, S. T., Maier, B., Yoshida, H., Seddu, K., Elbaz, N., Czysz, C., Zuk, O., Stranger, B. E., Ner-Gaon, H., Shay, T. (2019) ImmGen report: sexual dimorphism in the immune system transcriptome. Nat Commun 10, 4295.

46. Bray, N. L., Pimentel, H., Melsted, P., Pachter, L. (2016) Near-optimal probabilistic RNA-seq quantification. Nat Biotechnol 34, 525–7.

47. Stephens, A. S., Stephens, S. R., Morrison, N. A. (2011) Internal control genes for quantitative RT-PCR expression analysis in mouse osteoblasts, osteoclasts and macrophages. BMC Res Notes 4, 410.

48. Hawley, C. A., Rojo, R., Raper, A., Sauter, K. A., Lisowski, Z. M., Grabert, K., Bain, C. C., Davis, G. M., Louwe, P. A., Ostrowski, M. C., Hume, D. A., Pridans, C., Jenkins, S. J. (2018) Csf1r-mApple Transgene Expression and Ligand Binding In Vivo Reveal Dynamics of CSF1R Expression within the Mononuclear Phagocyte System. J Immunol 200, 2209–2223.

49. Sasmono, R. T., Oceandy, D., Pollard, J. W., Tong, W., Pavli, P., Wainwright, B. J., Ostrowski, M. C., Himes, S. R., Hume, D. A. (2003) A macrophage colony-stimulating factor receptor-green fluorescent protein transgene is expressed throughout the mononuclear phagocyte system of the mouse. Blood 101, 1155–63.

50. Schyns, J., Bai, Q., Ruscitti, C., Radermecker, C., De Schepper, S., Chakarov, S., Farnir, F., Pirottin, D., Ginhoux, F., Boeckxstaens, G., Bureau, F., Marichal, T. (2019) Non-classical tissue monocytes and two functionally distinct populations of interstitial macrophages populate the mouse lung. Nat Commun 10, 3964.

51. Hume, D. A., Summers, K. M., Rehli, M. (2016) Transcriptional Regulation and Macrophage Differentiation. Microbiol Spectr 4.

52. Rojo, R., Pridans, C., Langlais, D., Hume, D. A. (2017) Transcriptional mechanisms that control expression of the macrophage colony-stimulating factor receptor locus. Clin Sci (Lond) 131, 2161–2182.

53. Jubb, A. W., Young, R. S., Hume, D. A., Bickmore, W. A. (2016) Enhancer Turnover Is Associated with a Divergent Transcriptional Response to Glucocorticoid in Mouse and Human Macrophages. J Immunol 196, 813–822.

54. Audesse, A. J., Dhakal, S., Hassell, L. A., Gardell, Z., Nemtsova, Y., Webb, A. E. (2019) FOXO3 directly regulates an autophagy network to functionally regulate proteostasis in adult neural stem cells. PLoS Genet 15, e1008097.

55. Chen, Y., Wu, J., Liang, G., Geng, G., Zhao, F., Yin, P., Nowsheen, S., Wu, C., Li, Y., Li, L., Kim, W., Zhou, Q., Huang, J., Liu, J., Zhang, C., Guo, G., Deng, M., Tu, X., Gao, X., Liu, Z., Chen, Y., Lou, Z., Luo, K., Yuan, J. (2020) CHK2-FOXK axis promotes transcriptional control of autophagy programs. Sci Adv 6, eaax5819.

56. Schaffner, I., Minakaki, G., Khan, M. A., Balta, E. A., Schlotzer-Schrehardt, U., Schwarz, T. J., Beckervordersandforth, R., Winner, B., Webb, A. E., DePinho, R. A., Paik, J., Wurst, W., Klucken, J., Lie, D. C. (2018) FoxO Function Is Essential for Maintenance of Autophagic Flux and Neuronal Morphogenesis in Adult Neurogenesis. Neuron 99, 1188–1203 e6.

57. Choi, S., You, S., Kim, D., Choi, S. Y., Kwon, H. M., Kim, H. S., Hwang, D., Park, Y. J., Cho, C. S., Kim, W. U. (2017) Transcription factor NFAT5 promotes macrophage survival in rheumatoid arthritis. J Clin Invest 127, 954–969.

58. Roberts, T. L., Idris, A., Dunn, J. A., Kelly, G. M., Burnton, C. M., Hodgson, S., Hardy, L. L., Garceau, V., Sweet, M. J., Ross, I. L., Hume, D. A., Stacey, K. J. (2009) HIN-200 proteins regulate caspase activation in response to foreign cytoplasmic DNA. Science 323, 1057–60.

59. Briard, B., Place, D. E., Kanneganti, T. D. (2020) DNA Sensing in the Innate Immune Response. Physiology (Bethesda) 35, 112–124.

60. Zhao, Y., Shi, J., Shi, X., Wang, Y., Wang, F., Shao, F. (2016) Genetic functions of the NAIP family of inflammasome receptors for bacterial ligands in mice. J Exp Med 213, 647–56.

61. Pridans, C., Raper, A., Davis, G. M., Alves, J., Sauter, K. A., Lefevre, L., Regan, T., Meek, S., Sutherland, L., Thomson, A. J., Clohisey, S., Bush, S. J., Rojo, R., Lisowski, Z. M., Wallace, R., Grabert, K., Upton, K. R., Tsai, Y. T., Brown, D., Smith, L. B., Summers, K. M., Mabbott, N. A., Piccardo, P., Cheeseman, M. T., Burdon, T., Hume, D. A. (2018) Pleiotropic Impacts of Macrophage and Microglial Deficiency on Development in Rats with Targeted Mutation of the Csf1r Locus. J Immunol 201, 2683–2699.

62. Giotti, B., Chen, S. H., Barnett, M. W., Regan, T., Ly, T., Wiemann, S., Hume, D. A., Freeman, T. C. (2019) Assembly of a parts list of the human mitotic cell cycle machinery. J Mol Cell Biol 11, 703–718.

63. Aoki, M., Aoki, H., Ramanathan, R., Hait, N. C., Takabe, K. (2016) Sphingosine-1-Phosphate Signaling in Immune Cells and Inflammation: Roles and Therapeutic Potential. Mediators Inflamm 2016, 8606878.

64. Weichand, B., Popp, R., Dziumbla, S., Mora, J., Strack, E., Elwakeel, E., Frank, A. C., Scholich, K., Pierre, S., Syed, S. N., Olesch, C., Ringleb, J., Oren, B., Doring, C., Savai, R., Jung, M., von Knethen, A., Levkau, B., Fleming, I., Weigert, A., Brune, B. (2017) S1PR1 on tumor-associated macrophages promotes lymphangiogenesis and metastasis via NLRP3/IL-1beta. J Exp Med 214, 2695–2713.

65. Dutta, B., Arya, R. K., Goswami, R., Alharbi, M. O., Sharma, S., Rahaman, S. O. (2020) Role of macrophage TRPV4 in inflammation. Lab Invest 100, 178–185.

66. Morty, R. E. and Kuebler, W. M. (2014) TRPV4: an exciting new target to promote alveolocapillary barrier function. Am J Physiol Lung Cell Mol Physiol 307, L817–21.

67. Issitt, T., Bosseboeuf, E., De Winter, N., Dufton, N., Gestri, G., Senatore, V., Chikh, A., Randi, A. M., Raimondi, C. (2019) Neuropilin-1 Controls Endothelial Homeostasis by Regulating Mitochondrial Function and Iron-Dependent Oxidative Stress. iScience 11, 205–223.

68. Luo, B., Gan, W., Liu, Z., Shen, Z., Wang, J., Shi, R., Liu, Y., Liu, Y., Jiang, M., Zhang, Z., Wu, Y. (2016) Erythropoeitin Signaling in Macrophages Promotes Dying Cell Clearance and Immune Tolerance. Immunity 44, 287–302.

69. Hardbower, D. M., Singh, K., Asim, M., Verriere, T. G., Olivares-Villagomez, D., Barry, D. P., Allaman, M. M., Washington, M. K., Peek, R. M., Jr., Piazuelo, M. B., Wilson, K. T. (2016) EGFR regulates macrophage activation and function in bacterial infection. J Clin Invest 126, 3296–312.

70. Ivanov, S. and Randolph, G. J. (2017) Myeloid cells pave the way for lymphatic system development and maintenance. Pflugers Arch 469, 465–472.

71. Sanin, D. E., Matsushita, M., Klein Geltink, R. I., Grzes, K. M., van Teijlingen Bakker, N., Corrado, M., Kabat, A. M., Buck, M. D., Qiu, J., Lawless, S. J., Cameron, A. M., Villa, M., Baixauli, F., Patterson, A. E., Hassler, F., Curtis, J. D., O’Neill, C. M., O’Sullivan, D., Wu, D., Mittler, G., Huang, S. C., Pearce, E. L., Pearce, E. J. (2018) Mitochondrial Membrane Potential Regulates Nuclear Gene Expression in Macrophages Exposed to Prostaglandin E2. Immunity 49, 1021–1033 e6.

72. Yu, J., Zanotti, S., Schilling, L., Canalis, E. (2018) Nuclear factor of activated T cells 2 is required for osteoclast differentiation and function in vitro but not in vivo. J Cell Biochem 119, 9334–9345.

73. Van den Bossche, J., Malissen, B., Mantovani, A., De Baetselier, P., Van Ginderachter, J. A. (2012) Regulation and function of the E-cadherin/catenin complex in cells of the monocytemacrophage lineage and DCs. Blood 119, 1623–33.

74. Murray, P. J., Allen, J. E., Biswas, S. K., Fisher, E. A., Gilroy, D. W., Goerdt, S., Gordon, S., Hamilton, J. A., Ivashkiv, L. B., Lawrence, T., Locati, M., Mantovani, A., Martinez, F. O., Mege, J. L., Mosser, D. M., Natoli, G., Saeij, J. P., Schultze, J. L., Shirey, K. A., Sica, A., Suttles, J., Udalova, I., van Ginderachter, J. A., Vogel, S. N., Wynn, T. A. (2014) Macrophage activation and polarization: nomenclature and experimental guidelines. Immunity 41, 14–20.

75. Spadaro, O., Camell, C. D., Bosurgi, L., Nguyen, K. Y., Youm, Y. H., Rothlin, C. V., Dixit, V. D. (2017) IGF1 Shapes Macrophage Activation in Response to Immunometabolic Challenge. Cell Rep 19, 225–234.

76. Maridas, D. E., DeMambro, V. E., Le, P. T., Mohan, S., Rosen, C. J. (2017) IGFBP4 Is Required for Adipogenesis and Influences the Distribution of Adipose Depots. Endocrinology 158, 3488–3500.

77. Maridas, D. E., DeMambro, V. E., Le, P. T., Nagano, K., Baron, R., Mohan, S., Rosen, C. J. (2017) IGFBP-4 regulates adult skeletal growth in a sex-specific manner. J Endocrinol 233, 131–144.

78. Hume, D. A. (2006) The mononuclear phagocyte system. Curr Opin Immunol 18, 49–53.

79. Schulz, C., Gomez Perdiguero, E., Chorro, L., Szabo-Rogers, H., Cagnard, N., Kierdorf, K., Prinz, M., Wu, B., Jacobsen, S. E., Pollard, J. W., Frampton, J., Liu, K. J., Geissmann, F. (2012) A lineage of myeloid cells independent of Myb and hematopoietic stem cells. Science 336, 86–90.

80. Bain, C. C. and Schridde, A. (2018) Origin, Differentiation, and Function of Intestinal Macrophages. Front Immunol 9, 2733.

81. Briseno, C. G., Haldar, M., Kretzer, N. M., Wu, X., Theisen, D. J., Kc, W., Durai, V., Grajales-Reyes, G. E., Iwata, A., Bagadia, P., Murphy, T. L., Murphy, K. M. (2016) Distinct Transcriptional Programs Control Cross-Priming in Classical and Monocyte-Derived Dendritic Cells. Cell Rep 15, 2462–74.

82. Waddell, L. A., Lefevre, L., Bush, S. J., Raper, A., Young, R., Lisowski, Z. M., McCulloch, M. E. B., Muriuki, C., Sauter, K. A., Clark, E. L., Irvine, K. M., Pridans, C., Hope, J. C., Hume, D. A. (2018) ADGRE1 (EMR1, F4/80) Is a Rapidly-Evolving Gene Expressed in Mammalian Monocyte-Macrophages. Front Immunol 9, 2246.

83. Scott, C. L. and Guilliams, M. (2018) The role of Kupffer cells in hepatic iron and lipid metabolism. J Hepatol 69, 1197–1199.

84. Ma, F., Liu, S. Y., Razani, B., Arora, N., Li, B., Kagechika, H., Tontonoz, P., Nunez, V., Ricote, M., Cheng, G. (2014) Retinoid X receptor alpha attenuates host antiviral response by suppressing type I interferon. Nat Commun 5, 5494.

85. van der Spek, A. H., Fliers, E., Boelen, A. (2017) Thyroid hormone metabolism in innate immune cells. J Endocrinol 232, R67–R81.

86. Roussel-Gervais, A., Naciri, I., Kirsh, O., Kasprzyk, L., Velasco, G., Grillo, G., Dubus, P., Defossez, P. A. (2017) Loss of the Methyl-CpG-Binding Protein ZBTB4 Alters Mitotic Checkpoint, Increases Aneuploidy, and Promotes Tumorigenesis. Cancer Res 77, 62–73.

87. Haldar, M., Kohyama, M., So, A. Y., Kc, W., Wu, X., Briseno, C. G., Satpathy, A. T., Kretzer, N. M., Arase, H., Rajasekaran, N. S., Wang, L., Egawa, T., Igarashi, K., Baltimore, D., Murphy, T. L., Murphy, K. M. (2014) Heme-mediated SPI-C induction promotes monocyte differentiation into iron-recycling macrophages. Cell 156, 1223–1234.

88. Kohyama, M., Ise, W., Edelson, B. T., Wilker, P. R., Hildner, K., Mejia, C., Frazier, W. A., Murphy, T. L., Murphy, K. M. (2009) Role for Spi-C in the development of red pulp macrophages and splenic iron homeostasis. Nature 457, 318–21.

89. Bain, C. C., Hawley, C. A., Garner, H., Scott, C. L., Schridde, A., Steers, N. J., Mack, M., Joshi, A., Guilliams, M., Mowat, A. M., Geissmann, F., Jenkins, S. J. (2016) Long-lived selfrenewing bone marrow-derived macrophages displace embryo-derived cells to inhabit adult serous cavities. Nat Commun 7, comms11852.

90. Okabe, Y. and Medzhitov, R. (2014) Tissue-specific signals control reversible program of localization and functional polarization of macrophages. Cell 157, 832–44.

91. Costelloe, E. O., Stacey, K. J., Antalis, T. M., Hume, D. A. (1999) Regulation of the plasminogen activator inhibitor-2 (PAI-2) gene in murine macrophages. Demonstration of a novel pattern of responsiveness to bacterial endotoxin. J Leukoc Biol 66, 172–82.

92. Schroder, W. A., Hirata, T. D., Le, T. T., Gardner, J., Boyle, G. M., Ellis, J., Nakayama, E., Pathirana, D., Nakaya, H. I., Suhrbier, A. (2019) SerpinB2 inhibits migration and promotes a resolution phase signature in large peritoneal macrophages. Sci Rep 9, 12421.

93. Capucha, T., Mizraji, G., Segev, H., Blecher-Gonen, R., Winter, D., Khalaileh, A., Tabib, Y., Attal, T., Nassar, M., Zelentsova, K., Kisos, H., Zenke, M., Sere, K., Hieronymus, T., Burstyn-Cohen, T., Amit, I., Wilensky, A., Hovav, A. H. (2015) Distinct Murine Mucosal Langerhans Cell Subsets Develop from Pre-dendritic Cells and Monocytes. Immunity 43, 369–81.

94. Doebel, T., Voisin, B., Nagao, K. (2017) Langerhans Cells - The Macrophage in Dendritic Cell Clothing. Trends Immunol 38, 817–828.

95. Gross-Vered, M., Trzebanski, S., Shemer, A., Bernshtein, B., Curato, C., Stelzer, G., Salame, T. M., David, E., Boura-Halfon, S., Chappell-Maor, L., Leshkowitz, D., Jung, S. (2020) Defining murine monocyte differentiation into colonic and ileal macrophages. Elife 9. e49998

96. Hung, L. Y., Johnson, J. L., Ji, Y., Christian, D. A., Herbine, K. R., Pastore, C. F., Herbert, D. R. (2019) Cell-Intrinsic Wnt4 Influences Conventional Dendritic Cell Fate Determination to Suppress Type 2 Immunity. J Immunol 203, 511–519.

97. Sehgal, A., Donaldson, D. S., Pridans, C., Sauter, K. A., Hume, D. A., Mabbott, N. A. (2018) The role of CSF1R-dependent macrophages in control of the intestinal stem-cell niche. Nat Commun 9, 1272.

98. Shang, Y., Coppo, M., He, T., Ning, F., Yu, L., Kang, L., Zhang, B., Ju, C., Qiao, Y., Zhao, B., Gessler, M., Rogatsky, I., Hu, X. (2016) The transcriptional repressor Hes1 attenuates inflammation by regulating transcription elongation. Nat Immunol 17, 930–7.

99. Zhang, X., Li, X., Ning, F., Shang, Y., Hu, X. (2019) TLE4 acts as a corepressor of Hes1 to inhibit inflammatory responses in macrophages. Protein Cell 10, 300–305.

100. Schridde, A., Bain, C. C., Mayer, J. U., Montgomery, J., Pollet, E., Denecke, B., Milling, S. W. F., Jenkins, S. J., Dalod, M., Henri, S., Malissen, B., Pabst, O., McL Mowat, A. (2017) Tissuespecific differentiation of colonic macrophages requires TGFbeta receptor-mediated signaling. Mucosal Immunol 10, 1387–1399.

101. Morris, D. L., Singer, K., Lumeng, C. N. (2011) Adipose tissue macrophages: phenotypic plasticity and diversity in lean and obese states. Curr Opin Clin Nutr Metab Care 14, 341–6.

102. Lee, M. R., Lim, C. J., Lee, Y. H., Park, J. G., Sonn, S. K., Lee, M. N., Jung, I. H., Jeong, S. J., Jeon, S., Lee, M., Oh, K. S., Yang, Y., Kim, J. B., Choi, H. S., Jeong, W., Jeong, T. S., Yoon, W. K., Kim, H. C., Choi, J. H., Oh, G. T. (2014) The adipokine Retnla modulates cholesterol homeostasis in hyperlipidemic mice. Nat Commun 5, 4410.

103. Kumamoto, Y., Camporez, J. P. G., Jurczak, M. J., Shanabrough, M., Horvath, T., Shulman, G. I., Iwasaki, A. (2016) CD301b(+) Mononuclear Phagocytes Maintain Positive Energy Balance through Secretion of Resistin-like Molecule Alpha. Immunity 45, 583–596.

104. Fujita, K., Chakarov, S., Kobayashi, T., Sakamoto, K., Voisin, B., Duan, K., Nakagawa, T., Horiuchi, K., Amagai, M., Ginhoux, F., Nagao, K. (2019) Cell-autonomous FLT3L shedding via ADAM10 mediates conventional dendritic cell development in mouse spleen. Proc Natl Acad Sci U S A 116, 14714–14723.

105. Pool, L., Rivollier, A., Agace, W. W. (2020) Deletion of IRF4 in Dendritic Cells Leads to Delayed Onset of T Cell-Dependent Colitis. J Immunol 204, 1047–1055.

106. Anderson, D. A., 3rd, Murphy, K. M., Briseno, C. G. (2018) Development, Diversity, and Function of Dendritic Cells in Mouse and Human. Cold Spring Harb Perspect Biol 10.a028613

107. Forster, R., Davalos-Misslitz, A. C., Rot, A. (2008) CCR7 and its ligands: balancing immunity and tolerance. Nat Rev Immunol 8, 362–71.

108. Hildner, K., Edelson, B. T., Purtha, W. E., Diamond, M., Matsushita, H., Kohyama, M., Calderon, B., Schraml, B. U., Unanue, E. R., Diamond, M. S., Schreiber, R. D., Murphy, T. L., Murphy, K. M. (2008) Batf3 deficiency reveals a critical role for CD8alpha+ dendritic cells in cytotoxic T cell immunity. Science 322, 1097–100.

109. Bagadia, P., Huang, X., Liu, T. T., Durai, V., Grajales-Reyes, G. E., Nitschke, M., Modrusan, Z., Granja, J. M., Satpathy, A. T., Briseno, C. G., Gargaro, M., Iwata, A., Kim, S., Chang, H. Y., Shaw, A. S., Murphy, T. L., Murphy, K. M. (2019) An Nfil3-Zeb2-Id2 pathway imposes Irf8 enhancer switching during cDC1 development. Nat Immunol 20, 1174–1185.

110. Satpathy, A. T., Kc, W., Albring, J. C., Edelson, B. T., Kretzer, N. M., Bhattacharya, D., Murphy, T. L., Murphy, K. M. (2012) Zbtb46 expression distinguishes classical dendritic cells and their committed progenitors from other immune lineages. J Exp Med 209, 1135–52.

111. Salei, N., Rambichler, S., Salvermoser, J., Papaioannou, N. E., Schuchert, R., Pakalniskyte, D., Li, N., Marschner, J. A., Lichtnekert, J., Stremmel, C., Cernilogar, F. M., Salvermoser, M., Walzog, B., Straub, T., Schotta, G., Anders, H. J., Schulz, C., Schraml, B. U. (2020) The Kidney Contains Ontogenetically Distinct Dendritic Cell and Macrophage Subtypes throughout Development That Differ in Their Inflammatory Properties. J Am Soc Nephrol 31, 257–278.

112. Nagae, M., Ikeda, A., Hanashima, S., Kojima, T., Matsumoto, N., Yamamoto, K., Yamaguchi, Y. (2016) Crystal structure of human dendritic cell inhibitory receptor C-type lectin domain reveals the binding mode with N-glycan. FEBS Lett 590, 1280–8.

113. Troegeler, A., Mercier, I., Cougoule, C., Pietretti, D., Colom, A., Duval, C., Vu Manh, T. P., Capilla, F., Poincloux, R., Pingris, K., Nigou, J., Rademann, J., Dalod, M., Verreck, F. A., Al Saati, T., Lugo-Villarino, G., Lepenies, B., Hudrisier, D., Neyrolles, O. (2017) C-type lectin receptor DCIR modulates immunity to tuberculosis by sustaining type I interferon signaling in dendritic cells. Proc Natl Acad Sci U S A 114, E540–E549.

114. Uto, T., Fukaya, T., Takagi, H., Arimura, K., Nakamura, T., Kojima, N., Malissen, B., Sato, K. (2016) Clec4A4 is a regulatory receptor for dendritic cells that impairs inflammation and T-cell immunity. Nat Commun 7, 11273.

115. Wells, C. A., Salvage-Jones, J. A., Li, X., Hitchens, K., Butcher, S., Murray, R. Z., Beckhouse, A. G., Lo, Y. L., Manzanero, S., Cobbold, C., Schroder, K., Ma, B., Orr, S., Stewart, L., Lebus, D., Sobieszczuk, P., Hume, D. A., Stow, J., Blanchard, H., Ashman, R. B. (2008) The macrophage-inducible C-type lectin, mincle, is an essential component of the innate immune response to Candida albicans. J Immunol 180, 7404–13.

116. Link, V. M., Duttke, S. H., Chun, H. B., Holtman, I. R., Westin, E., Hoeksema, M. A., Abe, Y., Skola, D., Romanoski, C. E., Tao, J., Fonseca, G. J., Troutman, T. D., Spann, N. J., Strid, T., Sakai, M., Yu, M., Hu, R., Fang, R., Metzler, D., Ren, B., Glass, C. K. (2018) Analysis of Genetically Diverse Macrophages Reveals Local and Domain-wide Mechanisms that Control Transcription Factor Binding and Function. Cell 173, 1796–1809 e17.

117. Tamura, A., Hirai, H., Yokota, A., Kamio, N., Sato, A., Shoji, T., Kashiwagi, T., Torikoshi, Y., Miura, Y., Tenen, D. G., Maekawa, T. (2017) C/EBPbeta is required for survival of Ly6C(-) monocytes. Blood 130, 1809–1818.

118. Mildner, A., Schonheit, J., Giladi, A., David, E., Lara-Astiaso, D., Lorenzo-Vivas, E., Paul, F., Chappell-Maor, L., Priller, J., Leutz, A., Amit, I., Jung, S. (2017) Genomic Characterization of Murine Monocytes Reveals C/EBPbeta Transcription Factor Dependence of Ly6C(-) Cells. Immunity 46, 849–862 e7.

119. Bornstein, C., Winter, D., Barnett-Itzhaki, Z., David, E., Kadri, S., Garber, M., Amit, I. (2014) A negative feedback loop of transcription factors specifies alternative dendritic cell chromatin States. Mol Cell 56, 749–62.

120. Satpathy, A. T., Wu, X., Albring, J. C., Murphy, K. M. (2012) Re(de)fining the dendritic cell lineage. Nat Immunol 13, 1145–54.

121. Wu, X., Briseno, C. G., Durai, V., Albring, J. C., Haldar, M., Bagadia, P., Kim, K. W., Randolph, G. J., Murphy, T. L., Murphy, K. M. (2016) Mafb lineage tracing to distinguish macrophages from other immune lineages reveals dual identity of Langerhans cells. J Exp Med 213, 2553–2565.

122. Papadakis, K. A., Krempski, J., Svingen, P., Xiong, Y., Sarmento, O. F., Lomberk, G. A., Urrutia, R. A., Faubion, W. A. (2015) Kruppel-like factor KLF10 deficiency predisposes to colitis through colonic macrophage dysregulation. Am J Physiol Gastrointest Liver Physiol 309, G900–9.

123. van den Brink, S. C., Sage, F., Vertesy, A., Spanjaard, B., Peterson-Maduro, J., Baron, C. S., Robin, C., van Oudenaarden, A. (2017) Single-cell sequencing reveals dissociation-induced gene expression in tissue subpopulations. Nat Methods 14, 935–936.

124. Van Hove, H., Martens, L., Scheyltjens, I., De Vlaminck, K., Pombo Antunes, A. R., De Prijck, S., Vandamme, N., De Schepper, S., Van Isterdael, G., Scott, C. L., Aerts, J., Berx, G., Boeckxstaens, G. E., Vandenbroucke, R. E., Vereecke, L., Moechars, D., Guilliams, M., Van Ginderachter, J. A., Saeys, Y., Movahedi, K. (2019) A single-cell atlas of mouse brain macrophages reveals unique transcriptional identities shaped by ontogeny and tissue environment. Nat Neurosci 22, 1021–1035.

125. Pirzgalska, R. M., Seixas, E., Seidman, J. S., Link, V. M., Sanchez, N. M., Mahu, I., Mendes, R., Gres, V., Kubasova, N., Morris, I., Arus, B. A., Larabee, C. M., Vasques, M., Tortosa, F., Sousa, A. L., Anandan, S., Tranfield, E., Hahn, M. K., Iannacone, M., Spann, N. J., Glass, C. K., Domingos, A. I. (2017) Sympathetic neuron-associated macrophages contribute to obesity by importing and metabolizing norepinephrine. Nat Med 23, 1309–1318.

126. Ryan, D. G. and O’Neill, L. A. J. (2020) Krebs Cycle Reborn in Macrophage Immunometabolism. Annu Rev Immunol. Epub doi: 10.1146/annurev-immunol-081619-104850.

127. Gray, E. E., Friend, S., Suzuki, K., Phan, T. G., Cyster, J. G. (2012) Subcapsular sinus macrophage fragmentation and CD169+ bleb acquisition by closely associated IL-17-committed innate-like lymphocytes. PLoS One 7, e38258.

128. Lynch, R. W., Hawley, C. A., Pellicoro, A., Bain, C. C., Iredale, J. P., Jenkins, S. J. (2018) An efficient method to isolate Kupffer cells eliminating endothelial cell contamination and selective bias. J Leukoc Biol.104, 578–586

129. Bonnardel, J., T’Jonck, W., Gaublomme, D., Browaeys, R., Scott, C. L., Martens, L., Vanneste, B., De Prijck, S., Nedospasov, S. A., Kremer, A., Van Hamme, E., Borghgraef, P., Toussaint, W., De Bleser, P., Mannaerts, I., Beschin, A., van Grunsven, L. A., Lambrecht, B. N., Taghon, T., Lippens, S., Elewaut, D., Saeys, Y., Guilliams, M. (2019) Stellate Cells, Hepatocytes, and Endothelial Cells Imprint the Kupffer Cell Identity on Monocytes Colonizing the Liver Macrophage Niche. Immunity 51, 638–654 e9.

130. Ying, W., Lee, Y. S., Dong, Y., Seidman, J. S., Yang, M., Isaac, R., Seo, J. B., Yang, B. H., Wollam, J., Riopel, M., McNelis, J., Glass, C. K., Olefsky, J. M., Fu, W. (2019) Expansion of Islet-Resident Macrophages Leads to Inflammation Affecting beta Cell Proliferation and Function in Obesity. Cell Metab 29, 457–474 e5.

131. Li, W., Wang, Y., Zhao, H., Zhang, H., Xu, Y., Wang, S., Guo, X., Huang, Y., Zhang, S., Han, Y., Wu, X., Rice, C. M., Huang, G., Gallagher, P. G., Mendelson, A., Yazdanbakhsh, K., Liu, J., Chen, L., An, X. (2019) Identification and transcriptome analysis of erythroblastic island macrophages. Blood 134, 480–491.

132. A-Gonzales, N., Quintana, J. A., Garcia-Silva, S., Mazariegos, M., Gonzalez de la Aleja, A., Nicolas-Avila, J. A., Walter, W., Adrover, J. M., Crainiciuc, G., Kuchroo, V. K., Rothlin, C. V., Peinado, H., Castrillo, A., Ricote, M., Hidalgo, A. (2017) Phagocytosis imprints heterogeneity in tissue-resident macrophages. J Exp Med 214, 1281–1296.

133. Brown, C. C., Gudjonson, H., Pritykin, Y., Deep, D., Lavallee, V. P., Mendoza, A., Fromme, R., Mazutis, L., Ariyan, C., Leslie, C., Pe’er, D., Rudensky, A. Y. (2019) Transcriptional Basis of Mouse and Human Dendritic Cell Heterogeneity. Cell 179, 846–863 e24.

134. Lambert, S. A., Jolma, A., Campitelli, L. F., Das, P. K., Yin, Y., Albu, M., Chen, X., Taipale, J., Hughes, T. R., Weirauch, M. T. (2018) The Human Transcription Factors. Cell 175, 598–599.

135. Anderson, D. A., 3rd, Murphy, T. L., Eisenman, R. N., Murphy, K. M. (2020) The MYCL and MXD1 transcription factors regulate the fitness of murine dendritic cells. Proc Natl Acad Sci U S A.

136. Curi, R., de Siqueira Mendes, R., de Campos Crispin, L. A., Norata, G. D., Sampaio, S. C., Newsholme, P. (2017) A past and present overview of macrophage metabolism and functional outcomes. Clin Sci (Lond) 131, 1329–1342.

137. Liu, P. S., Wang, H., Li, X., Chao, T., Teav, T., Christen, S., Di Conza, G., Cheng, W. C., Chou, C. H., Vavakova, M., Muret, C., Debackere, K., Mazzone, M., Huang, H. D., Fendt, S. M., Ivanisevic, J., Ho, P. C. (2017) alpha-ketoglutarate orchestrates macrophage activation through metabolic and epigenetic reprogramming. Nat Immunol 18, 985–994.

138. Bhutia, Y. D. and Ganapathy, V. (2016) Glutamine transporters in mammalian cells and their functions in physiology and cancer. Biochim Biophys Acta 1863, 2531–9.

139. Freemerman, A. J., Zhao, L., Pingili, A. K., Teng, B., Cozzo, A. J., Fuller, A. M., Johnson, A. R., Milner, J. J., Lim, M. F., Galanko, J. A., Beck, M. A., Bear, J. E., Rotty, J. D., Bezavada, L., Smallwood, H. S., Puchowicz, M. A., Liu, J., Locasale, J. W., Lee, D. P., Bennett, B. J., Abel, E. D., Rathmell, J. C., Makowski, L. (2019) Myeloid Slc2a1-Deficient Murine Model Revealed Macrophage Activation and Metabolic Phenotype Are Fueled by GLUT1. J Immunol 202, 1265–1286.

140. Fang, H. Y., Hughes, R., Murdoch, C., Coffelt, S. B., Biswas, S. K., Harris, A. L., Johnson, R. S., Imityaz, H. Z., Simon, M. C., Fredlund, E., Greten, F. R., Rius, J., Lewis, C. E. (2009) Hypoxia-inducible factors 1 and 2 are important transcriptional effectors in primary macrophages experiencing hypoxia. Blood 114, 844–59.

141. Johnson, A. R., Qin, Y., Cozzo, A. J., Freemerman, A. J., Huang, M. J., Zhao, L., Sampey, B. P., Milner, J. J., Beck, M. A., Damania, B., Rashid, N., Galanko, J. A., Lee, D. P., Edin, M. L., Zeldin, D. C., Fueger, P. T., Dietz, B., Stahl, A., Wu, Y., Mohlke, K. L., Makowski, L. (2016) Metabolic reprogramming through fatty acid transport protein 1 (FATP1) regulates macrophage inflammatory potential and adipose inflammation. Mol Metab 5, 506–526.

142. Zhao, L., Cozzo, A. J., Johnson, A. R., Christensen, T., Freemerman, A. J., Bear, J. E., Rotty, J. D., Bennett, B. J., Makowski, L. (2017) Lack of myeloid Fatp1 increases atherosclerotic lesion size in Ldlr(-/-) mice. Atherosclerosis 266, 182–189.

143. Nomura, N., Verdon, G., Kang, H. J., Shimamura, T., Nomura, Y., Sonoda, Y., Hussien, S. A., Qureshi, A. A., Coincon, M., Sato, Y., Abe, H., Nakada-Nakura, Y., Hino, T., Arakawa, T., Kusano-Arai, O., Iwanari, H., Murata, T., Kobayashi, T., Hamakubo, T., Kasahara, M., Iwata, S., Drew, D. (2015) Structure and mechanism of the mammalian fructose transporter GLUT5. Nature 526, 397–401.

144. Caruana, B. T., Byrne, F. L., Knights, A. J., Quinlan, K. G. R., Hoehn, K. L. (2019) Characterization of Glucose Transporter 6 in Lipopolysaccharide-Induced Bone Marrow-Derived Macrophage Function. J Immunol 202, 1826–1832.

145. Lam-Yuk-Tseung, S., Picard, V., Gros, P. (2006) Identification of a tyrosine-based motif (YGSI) in the amino terminus of Nramp1 (Slc11a1) that is important for lysosomal targeting. J Biol Chem 281, 31677–88.

146. Wang, L., Fang, B., Fujiwara, T., Krager, K., Gorantla, A., Li, C., Feng, J. Q., Jennings, M. L., Zhou, J., Aykin-Burns, N., Zhao, H. (2018) Deletion of ferroportin in murine myeloid cells increases iron accumulation and stimulates osteoclastogenesis in vitro and in vivo. J Biol Chem 293, 9248–9264.

147. Kapetanovic, R., Bokil, N. J., Achard, M. E., Ong, C. L., Peters, K. M., Stocks, C. J., Phan, M. D., Monteleone, M., Schroder, K., Irvine, K. M., Saunders, B. M., Walker, M. J., Stacey, K. J., McEwan, A. G., Schembri, M. A., Sweet, M. J. (2016) Salmonella employs multiple mechanisms to subvert the TLR-inducible zinc-mediated antimicrobial response of human macrophages. FASEB J 30, 1901–12.

148. Stafford, S. L., Bokil, N. J., Achard, M. E., Kapetanovic, R., Schembri, M. A., McEwan, A. G., Sweet, M. J. (2013) Metal ions in macrophage antimicrobial pathways: emerging roles for zinc and copper. Biosci Rep 33.

149. Xu, H., Ghishan, F. K., Kiela, P. R. (2018) SLC9 Gene Family: Function, Expression, and Regulation. Compr Physiol 8, 555–583.

150. Hume, D. A. and Gordon, S. (1983) Mononuclear phagocyte system of the mouse defined by immunohistochemical localization of antigen F4/80. Identification of resident macrophages in renal medullary and cortical interstitium and the juxtaglomerular complex. J Exp Med 157, 1704–9.

151. Viehmann, S. F., Bohner, A. M. C., Kurts, C., Brahler, S. (2018) The multifaceted role of the renal mononuclear phagocyte system. Cell Immunol.

152. Puranik, A. S., Leaf, I. A., Jensen, M. A., Hedayat, A. F., Saad, A., Kim, K. W., Saadalla, A. M., Woollard, J. R., Kashyap, S., Textor, S. C., Grande, J. P., Lerman, A., Simari, R. D., Randolph, G. J., Duffield, J. S., Lerman, L. O. (2018) Kidney-resident macrophages promote a proangiogenic environment in the normal and chronically ischemic mouse kidney. Sci Rep 8, 13948.

153. Stamatiades, E. G., Tremblay, M. E., Bohm, M., Crozet, L., Bisht, K., Kao, D., Coelho, C., Fan, X., Yewdell, W. T., Davidson, A., Heeger, P. S., Diebold, S., Nimmerjahn, F., Geissmann, F. (2016) Immune Monitoring of Trans-endothelial Transport by Kidney-Resident Macrophages. Cell 166, 991–1003.

154. Lee, A. S., Lee, J. E., Jung, Y. J., Kim, D. H., Kang, K. P., Lee, S., Park, S. K., Lee, S. Y., Kang, M. J., Moon, W. S., Kim, H. J., Jeong, Y. B., Sung, M. J., Kim, W. (2013) Vascular endothelial growth factor-C and -D are involved in lymphangiogenesis in mouse unilateral ureteral obstruction. Kidney Int 83, 50–62.

155. Himes, S. R., Cronau, S., Mulford, C., Hume, D. A. (2005) The Runx1 transcription factor controls CSF-1-dependent and -independent growth and survival of macrophages. Oncogene 24, 5278–86.

156. Ginhoux, F., Greter, M., Leboeuf, M., Nandi, S., See, P., Gokhan, S., Mehler, M. F., Conway, S. J., Ng, L. G., Stanley, E. R., Samokhvalov, I. M., Merad, M. (2010) Fate mapping analysis reveals that adult microglia derive from primitive macrophages. Science 330, 841–5.

157. Giladi, A. and Amit, I. (2018) Single-Cell Genomics: A Stepping Stone for Future Immunology Discoveries. Cell 172, 14–21.

158. Gunther, P. and Schultze, J. L. (2019) Mind the Map: Technology Shapes the Myeloid Cell Space. Front Immunol 10, 2287.

159. Andrews, T. S. and Hemberg, M. (2018) Identifying cell populations with scRNASeq. Mol Aspects Med 59, 114–122.

160. Chen, G., Ning, B., Shi, T. (2019) Single-Cell RNA-Seq Technologies and Related Computational Data Analysis. Front Genet 10, 317.

161. Becht, E., McInnes, L., Healy, J., Dutertre, C. A., Kwok, I. W. H., Ng, L. G., Ginhoux, F., Newell, E. W. (2018) Dimensionality reduction for visualizing single-cell data using UMAP. Nat Biotechnol. Epub doi: 10.1038/nbt.4314.

162. Aitchison, L., Corradi, N., Latham, P. E. (2016) Zipf’s Law Arises Naturally When There Are Underlying, Unobserved Variables. PLoS Comput Biol 12, e1005110.

163. Ueda, H. R., Hayashi, S., Matsuyama, S., Yomo, T., Hashimoto, S., Kay, S. A., Hogenesch, J. B., Iino, M. (2004) Universality and flexibility in gene expression from bacteria to human. Proc Natl Acad Sci U S A 101, 3765–9.

164. Gibbings, S. L., Thomas, S. M., Atif, S. M., McCubbrey, A. L., Desch, A. N., Danhorn, T., Leach, S. M., Bratton, D. L., Henson, P. M., Janssen, W. J., Jakubzick, C. V. (2017) Three Unique Interstitial Macrophages in the Murine Lung at Steady State. Am J Respir Cell Mol Biol 57, 66–76.

165. Tan, S. Y. and Krasnow, M. A. (2016) Developmental origin of lung macrophage diversity. Development 143, 1318–27.

166. Hume, D. A. (2000) Probability in transcriptional regulation and its implications for leukocyte differentiation and inducible gene expression. Blood 96, 2323–8.

167. Reinius, B., Mold, J. E., Ramskold, D., Deng, Q., Johnsson, P., Michaelsson, J., Frisen, J., Sandberg, R. (2016) Analysis of allelic expression patterns in clonal somatic cells by single-cell RNA-seq. Nat Genet 48, 1430–1435.

168. Zilionis, R., Engblom, C., Pfirschke, C., Savova, V., Zemmour, D., Saatcioglu, H. D., Krishnan, I., Maroni, G., Meyerovitz, C. V., Kerwin, C. M., Choi, S., Richards, W. G., De Rienzo, A., Tenen, D. G., Bueno, R., Levantini, E., Pittet, M. J., Klein, A. M. (2019) Single-Cell Transcriptomics of Human and Mouse Lung Cancers Reveals Conserved Myeloid Populations across Individuals and Species. Immunity 50, 1317–1334 e10.

169. van Vugt, M. J., Kleijmeer, M. J., Keler, T., Zeelenberg, I., van Dijk, M. A., Leusen, J. H., Geuze, H. J., van de Winkel, J. G. (1999) The FcgammaRIa (CD64) ligand binding chain triggers major histocompatibility complex class II antigen presentation independently of its associated FcR gamma-chain. Blood 94, 808–17.

170. Orecchioni, M., Ghosheh, Y., Pramod, A. B., Ley, K. (2019) Macrophage Polarization: Different Gene Signatures in M1(LPS+) vs. Classically and M2(LPS-) vs. Alternatively Activated Macrophages. Front Immunol 10, 1084.

171. Jenkins, S. J., Ruckerl, D., Thomas, G. D., Hewitson, J. P., Duncan, S., Brombacher, F., Maizels, R. M., Hume, D. A., Allen, J. E. (2013) IL-4 directly signals tissue-resident macrophages to proliferate beyond homeostatic levels controlled by CSF-1. J Exp Med 210, 2477–91.

172. Guilliams, M. and Scott, C. L. (2017) Does niche competition determine the origin of tissue-resident macrophages? Nat Rev Immunol 17, 451–460.

173. T’Jonck, W., Guilliams, M., Bonnardel, J. (2018) Niche signals and transcription factors involved in tissue-resident macrophage development. Cell Immunol.

174. Geissmann, F., Gordon, S., Hume, D. A., Mowat, A. M., Randolph, G. J. (2010) Unravelling mononuclear phagocyte heterogeneity. Nat Rev Immunol 10, 453–60.

175. Fujiu, K., Shibata, M., Nakayama, Y., Ogata, F., Matsumoto, S., Noshita, K., Iwami, S., Nakae, S., Komuro, I., Nagai, R., Manabe, I. (2017) A heart-brain-kidney network controls adaptation to cardiac stress through tissue macrophage activation. Nat Med 23, 611–622.

176. Wolf, Y., Boura-Halfon, S., Cortese, N., Haimon, Z., Sar Shalom, H., Kuperman, Y., Kalchenko, V., Brandis, A., David, E., Segal-Hayoun, Y., Chappell-Maor, L., Yaron, A., Jung, S. (2017) Brown-adipose-tissue macrophages control tissue innervation and homeostatic energy expenditure. Nat Immunol 18, 665–674.

177. Shemer, A., Grozovski, J., Tay, T. L., Tao, J., Volaski, A., Suss, P., Ardura-Fabregat, A., Gross-Vered, M., Kim, J. S., David, E., Chappell-Maor, L., Thielecke, L., Glass, C. K., Cornils, K., Prinz, M., Jung, S. (2018) Engrafted parenchymal brain macrophages differ from microglia in transcriptome, chromatin landscape and response to challenge. Nat Commun 9, 5206.

178. Li, Q., Cheng, Z., Zhou, L., Darmanis, S., Neff, N. F., Okamoto, J., Gulati, G., Bennett, M. L., Sun, L. O., Clarke, L. E., Marschallinger, J., Yu, G., Quake, S. R., Wyss-Coray, T., Barres, B. A. (2019) Developmental Heterogeneity of Microglia and Brain Myeloid Cells Revealed by Deep Single-Cell RNA Sequencing. Neuron 101, 207–223 e10.

179. Rauschmeier, R., Gustafsson, C., Reinhardt, A., N, A. G., Tortola, L., Cansever, D., Subramanian, S., Taneja, R., Rossner, M. J., Sieweke, M. H., Greter, M., Mansson, R., Busslinger, M., Kreslavsky, T. (2019) Bhlhe40 and Bhlhe41 transcription factors regulate alveolar macrophage self-renewal and identity. EMBO J 38, e101233.

180. Jaitin, D. A., Adlung, L., Thaiss, C. A., Weiner, A., Li, B., Descamps, H., Lundgren, P., Bleriot, C., Liu, Z., Deczkowska, A., Keren-Shaul, H., David, E., Zmora, N., Eldar, S. M., Lubezky, N., Shibolet, O., Hill, D. A., Lazar, M. A., Colonna, M., Ginhoux, F., Shapiro, H., Elinav, E., Amit, I. (2019) Lipid-Associated Macrophages Control Metabolic Homeostasis in a Trem2-Dependent Manner. Cell 178, 686–698 e14.

181. Thion, M. S., Low, D., Silvin, A., Chen, J., Grisel, P., Schulte-Schrepping, J., Blecher, R., Ulas, T., Squarzoni, P., Hoeffel, G., Coulpier, F., Siopi, E., David, F. S., Scholz, C., Shihui, F., Lum, J., Amoyo, A. A., Larbi, A., Poidinger, M., Buttgereit, A., Lledo, P. M., Greter, M., Chan, J. K. Y., Amit, I., Beyer, M., Schultze, J. L., Schlitzer, A., Pettersson, S., Ginhoux, F., Garel, S. (2018) Microbiome Influences Prenatal and Adult Microglia in a Sex-Specific Manner. Cell 172, 500–516 e16.

182. Stock, A. T., Collins, N., Smyth, G. K., Hu, Y., Hansen, J. A., D’Silva, D. B., Jama, H. A., Lew, A. M., Gebhardt, T., McLean, C. A., Wicks, I. P. (2019) The Selective Expansion and Targeted Accumulation of Bone Marrow-Derived Macrophages Drive Cardiac Vasculitis. J Immunol 202, 3282–3296.

